# SNARE disassembly requires Sec18/NSF side-loading

**DOI:** 10.1101/2024.08.30.610324

**Authors:** Yousuf A. Khan, K. Ian White, Richard A. Pfuetzner, Bharti Singal, Luis Esquivies, Garvey Mckenzie, Fang Liu, Katherine DeLong, Ucheor B. Choi, Elizabeth Montabana, Theresa Mclaughlin, William T. Wickner, Axel T. Brunger

## Abstract

SNARE proteins drive membrane fusion at different cell compartments as their core domains zipper into a parallel four-helix bundle. After fusion, these bundles are disassembled by the AAA+ protein Sec18/NSF and its adaptor Sec17/α-SNAP to make them available for subsequent rounds of membrane fusion. SNARE domains are often flanked by C-terminal transmembrane or N-terminal domains. Previous structures of the NSF–α-SNAP–SNARE complex revealed binding to the D1 ATPase pore, posing a topological constraint as SNARE transmembrane domains would prevent complete substrate threading as suggested for other AAA+ systems. Using mass-spectrometry in yeast cells, we show N-terminal SNARE domain interactions with Sec18, exacerbating this topological issue. We present cryo-EM structures of a yeast SNARE complex, Sec18, and Sec17 in a non-hydrolyzing condition, which show SNARE Sso1 threaded through the D1 and D2 ATPase rings of Sec18, with its folded, N-terminal Habc domain interacting with the D2 ring. This domain does not unfold during Sec18/NSF activity. Cryo-EM structures under hydrolyzing conditions revealed substrate-released and substrate-free states of Sec18 with a coordinated opening in the side of the ATPase rings. Thus, Sec18/NSF operates by substrate side-loading and unloading topologically constrained SNARE substrates.

## Main Text

Cellular compartmentalization, growth, hormone secretion, transport, neurotransmission, and many other pathways depend on precise, rapid, and regulated membrane fusion^1,2^. Membrane fusion in eukaryotic cells is mediated by a highly conserved superfamily of SNAREs (Soluble N-ethylmaleimide sensitive factor Attachment protein Receptors)^3^. All SNAREs share a characteristic 60–70 amino acid SNARE domain often flanked by a C-terminal transmembrane domain, membrane anchors, and a folded N-terminal variable domain specific to the SNARE’s function and intracellular pathway^4^. SNAREs on opposing membranes interact primarily through their SNARE domains to form a parallel *trans*-SNARE complex, juxtaposing two different membranes^5,6^. Membrane fusion commences when these SNARE domains zipper together in a directed fashion^7,8^. After fusion, the SNAREs form a highly stable parallel helical bundle, the so-called *cis*-SNARE complex^8^. This highly stable four-helix *cis*-SNARE complex is disassembled by Sec18 (N-ethylmaleimide-sensitive factor, NSF, in higher eukaryotes)^9–11^ to provide the energy for subsequent membrane fusion events^12^.

Sec18/NSF is a universally conserved AAA+ (ATPases associated with diverse cellular activities) protein translocase^13–15^. Sec18/NSF consists of an N-domain, an active D1 AAA+ domain, and a catalytically inactive D2 AAA+ oligomerization domain^16^. It was initially discovered as a critical complementation group required by the yeast secretory pathway, and it is now known for its role in recycling SNAREs^17^ and SNARE assembly quality control for proper SNARE complex assembly^18–21^. Sec18/NSF, with the adaptor protein Sec17 (α-SNAP in higher eukaryotes), disassembles *cis*-SNARE complexes. The disassembly requires multiple ATP hydrolysis events in the presence of Mg^2+^; as few as 6 ATP hydrolysis events are sufficient^22,23^.

Cryo-EM structures of the mammalian 20S complex (NSF, α-SNAP, and neuronal SNAREs ternary complex) in a non-hydrolyzing condition (*i.e.*, in the absence of divalent cations) revealed a supramolecular architecture in which the NSF N-domains bind α-SNAP molecules, three or four of which in turn bind the four-helix SNARE complex consisting of the SNARE domains of syntaxin-1A, SNAP-25, and synaptobrevin^10,24^. The N-terminal residues of the SNARE domain of SNAP-25 were bound to the D1 ring pore without apparent ATP hydrolysis, where it interacts with several conserved tyrosine amino acids in a spiral staircase-like pattern^10,13,25^. In these EM maps, no ordered density was observed in the D2 ring pore, consistent with the notion that it is catalytically inactive and primarily responsible for NSF oligomerization rather than substrate engagement^26^. The interaction between the SNARE substrate and the D1 pore is like that observed for other AAA+ translocases and suggests a conserved mechanism for substrate threading through the D1 pore. However, the membrane anchors and domains of the SNAREs would seemingly prevent complete threading of the type suggested for other AAA+ systems^27–29^, posing a topological challenge.

Furthermore, SNAREs often contain globular N-terminal domain(s) of variable length and structure; for example, the N-terminal domain of syntaxin consists of a three-helix bundle (Habc domain) involved in regulating its function^30^, and the N-terminal domains of Use1 and Sec20 in part form a stable 255 kDa tethering complex^31^. Complete threading would imply that such N-terminal domains are somehow unfolded. These topological constraints surrounding SNARE loading, processing, and release through Sec18/NSF are further compounded by the observation that Sec18/NSF disassembles all SNAREs in all cellular contexts^3,4,32^, which all contain a variety of different N- and C-terminal domains and membrane arrangements/linkages.

### *In vivo* cross-linking mass-spectrometry with Sec18

These questions led us to investigate the space of Sec18/NSF—SNARE interactions through *in vivo* protein crosslinking mass-spectrometry (XL-MS) in yeast. Due to its power as a model system and relatively simple SNARE proteome, *S. cerevisiae* is an excellent model for investigating Sec18 interactions with different substrates in live cells. As such, we developed a protocol for *in vivo* crosslinking mass-spectrometry (XL-MS) in yeast to identify binding partners of Sec18 (**Extended Data Figure 1A, Methods**).

In total, 35 identified proteins are shared between disuccinimidyl glutarate crosslinker (DSG) treated and untreated conditions (*i.e.*, they are non-specific proteins) (**Figure 1A**). The nine proteins observed only in the untreated condition had few unique spectra mapping to their identified proteins, suggesting that they were not detected in the treated condition due to their low abundance and difficulty of consistent detection. The 205 proteins unique to the 5 mM DSG treated condition were processed using spatial analysis of functional enrichment (SAFE) analysis to visualize the various cellular processes to which the identified proteins contribute^33^. At increasing levels of significance thresholds (*p =* 0.01, 0.001, and 0.0001), the identified proteins only enriched the vesicle trafficking processes. Gene ontology (GO) analysis also revealed lesser-known processes, such as vacuolar acidification and ergosterol biosynthesis (**Supplementary Figure 1**), consistent with Sec18’s functions in these contexts^34–36^. The enrichment of these processes validated our crosslinking protocol for specifically targeting Sec18 and its binding partners inside the cell.

**Figure 1:**
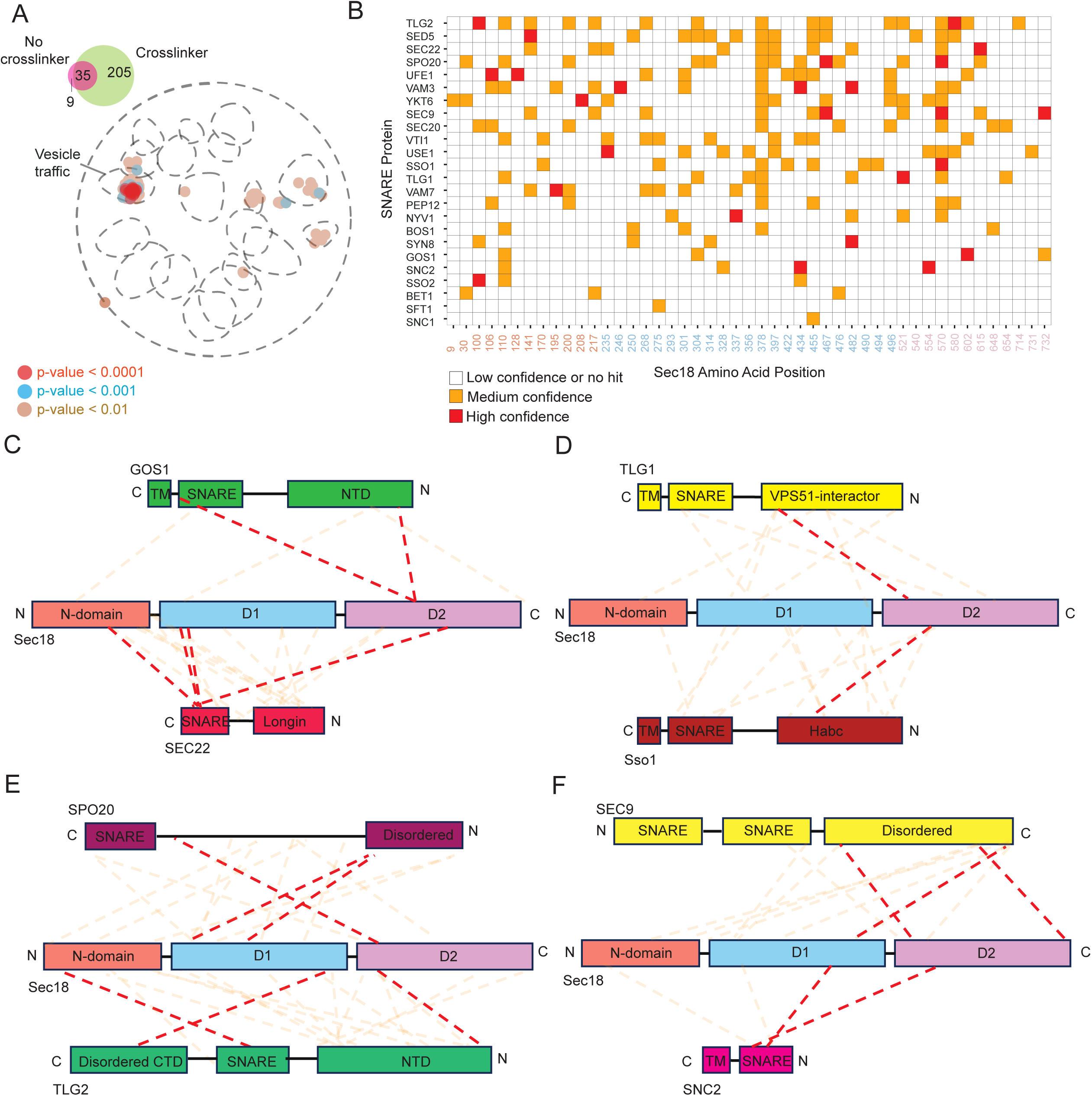
*In vivo* crosslinking mass-spectrometry of Sec18. **a,** Top left: Venn diagram of proteins found in the condition without a crosslinking agent (red) added and with crosslinking agent added (green). Bottom right: SAFE-analysis^33^ of yeast proteins that specifically crosslink to Sec18. Blobs represent enrichments in the pathway. Blobs of different colors represent enrichments that are specific to different p-value thresholds (brown < 0.01, blue < 0.001, red < 0.0001. p-values are derived from Fisher’s exact test with Bonferroni correction for multiple testing). **b,** Heatmap of yeast SNARE proteins crosslinking to specific Sec18 positions. The Sec18 primary sequence along the x-axis is colored coded by domain, and yeast SNARE proteins with at least one medium confidence crosslink are shown along the y-axis. Red squares represent positions where one or more manually verified, high-confidence crosslinks were found (**Methods**). Orange squares represent where one or more manually verified medium-confidence crosslinks were found. White squares represent positions in which manually verified crosslinks were not found or were of low confidence. **c-f,** Crosslinking schematic of Sec18 and SNAREs that contain at least one high-confidence crosslink to the Sec18 D2 domain. Red dashed lines represent high-confidence crosslinks, and orange dashed lines represent medium-confidence crosslinks.

Within our enriched protein dataset, we found many yeast SNAREs involved in different pathways and compartments of the cell. We then mapped crosslinked residues of these SNAREs to those of Sec18 (**Figure 1B**). Using a series of empirical constraints (**Methods**), we classified a crosslink as high confidence if it met all constraints, medium confidence if it met some but not all, and low confidence if it did not. Since crosslinking was performed in live cells, we expect these crosslinks to represent interactions between SNAREs and Sec18 during substrate loading, disassembly, and substrate release.

Previous cryo-EM structures of the complex of NSF, α-SNAP, and neuronal SNAREs in a non-hydrolyzing condition revealed that the four-helix SNARE bundle interacts with between two and four α-SNAP molecules, which in turn are bound by the N-domains of NSF; the N-terminal end of one of the SNAREs is bound to the pore of the D1 ring^10,24,25^. Consistent with these structures, we found crosslinks between SNARE domains and the N-domain or D1 domain (examples in **Figure 1C, E, F**). Additionally, we observed several high-confidence crosslinks of yeast SNARE proteins to the D2 domain, an unexpected result given an absence of SNARE density in the D2 ring in these previous cryo-EM structures and that the D2 domain is catalytically inactive^16^ and has had no previously reported role in SNARE recycling. Considering this unexpected result, we thus focused on SNAREs with at least one high-confidence crosslink to the D2 domain of Sec18 (**Figures 1C-F**).

Specifically, seven of ten high-confidence D2 crosslinks connect to regions N-terminal to a SNARE domain (**Figures 1C-F**). For example, for Sso1, the high-confidence crosslink to D2 involves its Habc domain (**Figure 1D, Extended Data Figure 1B**). The Habc domain is a stable three-helix bundle^30^; thus, assuming complete threading through the D1 and D2 pores, the Habc domain (∼30 Å diameter) would have to be unfolded transiently. The remaining three high-confidence D2 crosslinks connect to C-terminal regions within SNARE domains. At first glance, this suggests complete substrate threading through both the D1 and D2 rings, akin to other AAA+ protein translocases^37^. However, considering the topology imposed by membrane domains, anchors, and folded N-terminal domains, how is a SNARE substrate loaded and released? To answer these vexing topological questions, we next determined cryo-EM structures of this orthologous yeast complex together with Sec18 and Sec17.

### The Sec18—Sec17—Sso1—Snc1—Sec9 (y20S) complex

To corroborate the surprising LC-MS/MS results, we determined cryo-EM structures of a yeast SNARE complex together with Sec18 and Sec17. We chose the Sso1**—**Sec9 **—**Snc1/Snc2 complex (referred to as ySNARE complex) since each component of this complex crosslinked with high confidence to the D2 domain of Sec18 (Snc1 is highly homologous to Snc2). We prepared the Sec18—Sec17—Sso1—Snc1—Sec9 (y20S) complex in a non-hydrolyzing condition (**Extended Data Figure 2, Methods**), and after single particle cryo-EM data collection and processing, we obtained 381,591 high-quality particles that yielded eight 3D classes into which models were built (**Figure 2A–E, Supplementary Figure 2A-B, Supplementary Table 1**).

**Figure 2:**
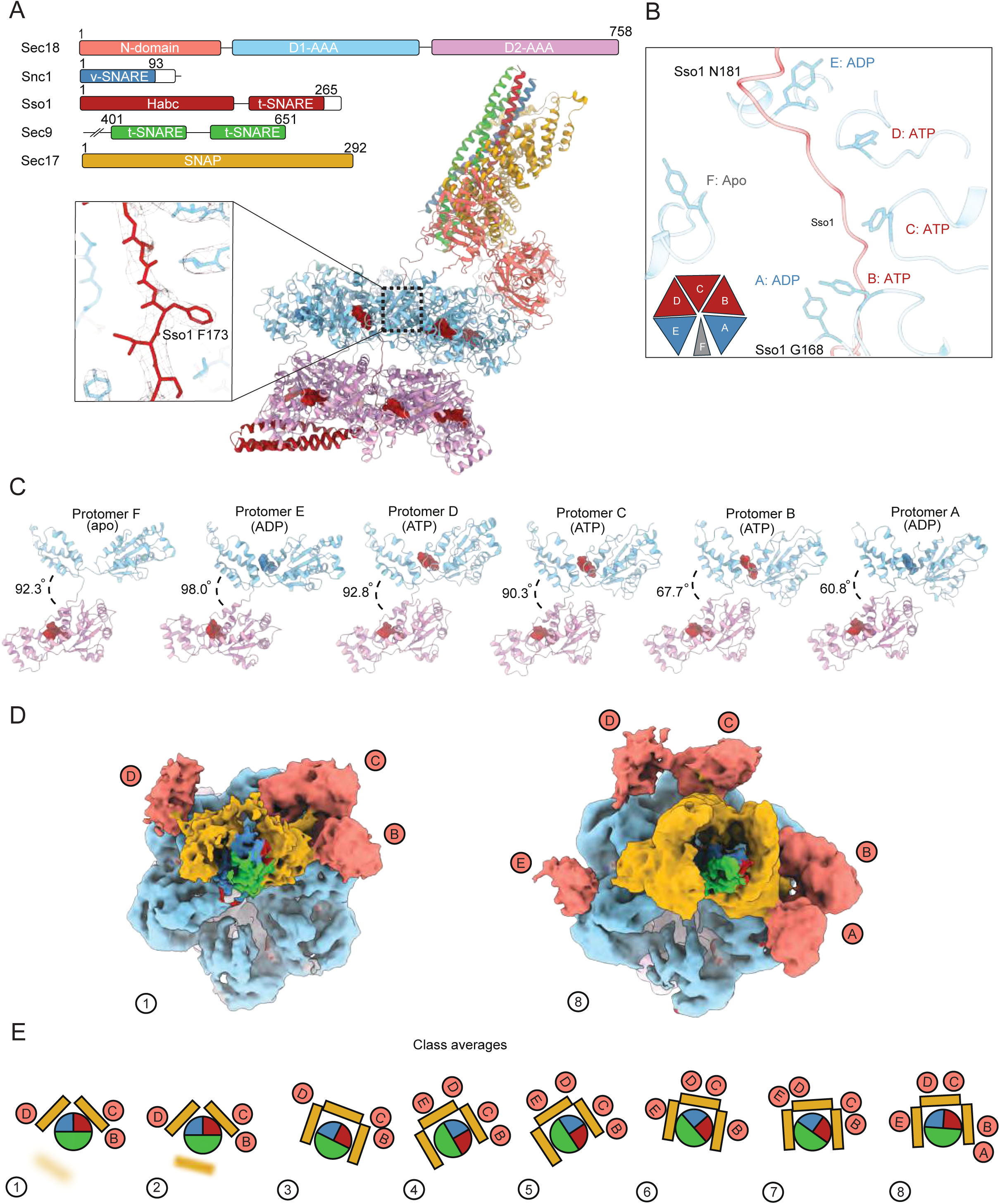
The y20S supramolecular complex. **a,** Domain diagrams of the components used to prepare the y20S complex; coloring follows this throughout (top left). The structure of class 1 of y20S, with ATP and ADP nucleotides (red and blue, respectively; bottom right). Cryo-EM density for class 1 and the Sso1 atomic model around residue F173 (inset, bottom left). **b,** Sso1 substrate is engaged by Y315 in all D1 protomers with bound nucleotide. **c,** Conformations of the protomers of Sec18 when engaged to substrate. **d**, Top-down views of class 1 and class 8 cryo-EM maps. **e,** Cartoon top-down views of eight y20S classes. These classes differ primarily by spire configuration, *i.e.*, the pattern of N-domain and ɑ-SNAP engagement.

In the structures of all eight classes, Sso1 is threaded through the D1 pore of Sec18 (**Figure 2A**). A characteristic phenylalanine side chain density in Sso1 is present at the same position relative to Sec18 in all classes, allowing for reliable indexing of Sso1. The nucleotide states and the arrangements of D1 around the substrate were also the same among all eight classes, with the ADP-engaged E protomer forming the top of a spiral staircase of tyrosine residues. Protomers D, B, and C were all ATP bound and formed the middle of the staircase. The bottom protomer, A, forms the base of the staircase. Protomer F, with no apparent density for nucleotide in D1, was not associated with Sso1 (**Figure 2B, Extended Data Figure 3A**). This arrangement of the protomers is driven, in part, by the angle between the more mobile D1 domain and rigid D2 domain. From protomers E to A, the angle between the D1 and D2 decreases monotonically (**Figure 2C**). The protomer at the top of the staircase maintains the largest angle between D1 and D2, while the one at the bottom has the smallest. The eight classes from the non-hydrolyzing y20S dataset also show differences in the ySNARE-Sec17-Sec18-N-domain subcomplex arrangement (“spire”) and Sec17 stoichiometry above the D1 ring of the y20S complex before disassembly (**Figure 2D-E**, **Figure 3A-B**; **Supplementary Discussion**).

**Figure 3:**
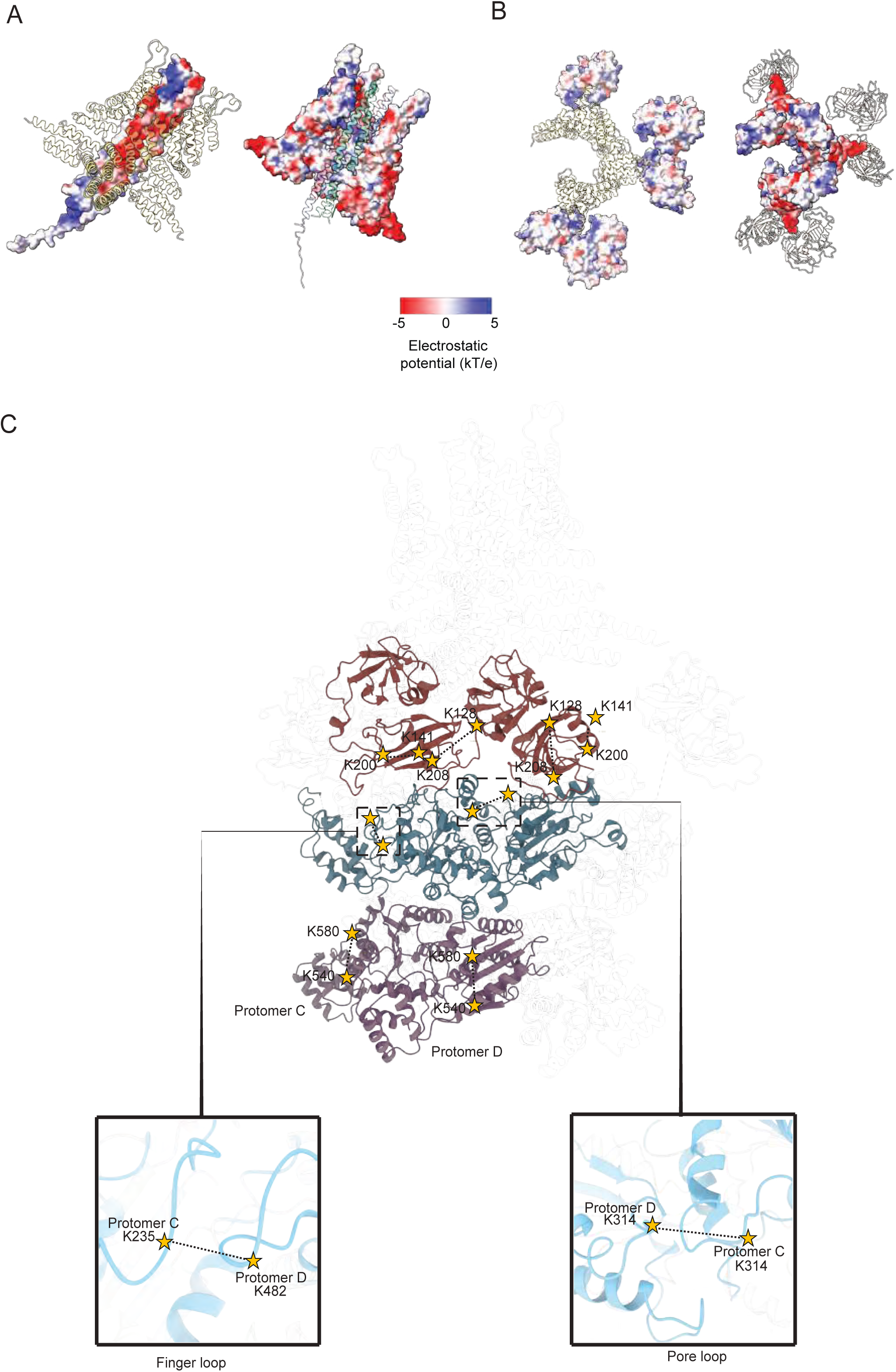
Intra and inter-protomer crosslinks mapped on the y20S complex structure. **a,** Electrostatic surface potentials of the interaction between SNAREs and Sec17. Negatively charged regions are colored red, and positive ones are colored blue. **b,** Electrostatic surface potentials of the interaction between Sec17 and the N-domains of Sec18. Alternating views of these interactions are shown to demonstrate the complementarity of the electrostatic interactions. **c,** For clarity, only two Sec18 protomers of the y20S complex are shown. The black dashed lines represent crosslinks designated as high quality and superimposed over the diagram. The two insets show close-up views of two inter-protomer crosslinks.

The high-confidence crosslinks for Sec18 intra- and inter-protomer interactions are consistent with the cryo-EM structure, specifically with the arrangements of the N-domain and D2-domain (**Figure 3C**, **Figure 4A**). The high-confidence crosslink between the D1 domain of Sec18 and Snc2 is also consistent with our cryo-EM structures: the D1 domain is proximal to the SNARE domain of Snc2 in the initial loading state of the 20S complex where Sso1 is loaded into the D1 pore. The other high-confidence crosslink occurs between Snc2 and a D2 domain residue at the interface between the D1 and D2 protomers. However, there is no density for Snc2 in our cryo-EM structures in the D1 pore, suggesting that these crosslinks may represent a transient state. Moreover, inter-protomer crosslinks were found between pore loop regions, and between the inter-protomer loop between residues 476 and 490 and N-domain linker regions of D1. These high-confidence crosslinks were within the theoretical maximum crosslinking distance (N_ζ_–N_ζ_: ∼40 Å)^38^.

**Figure 4:**
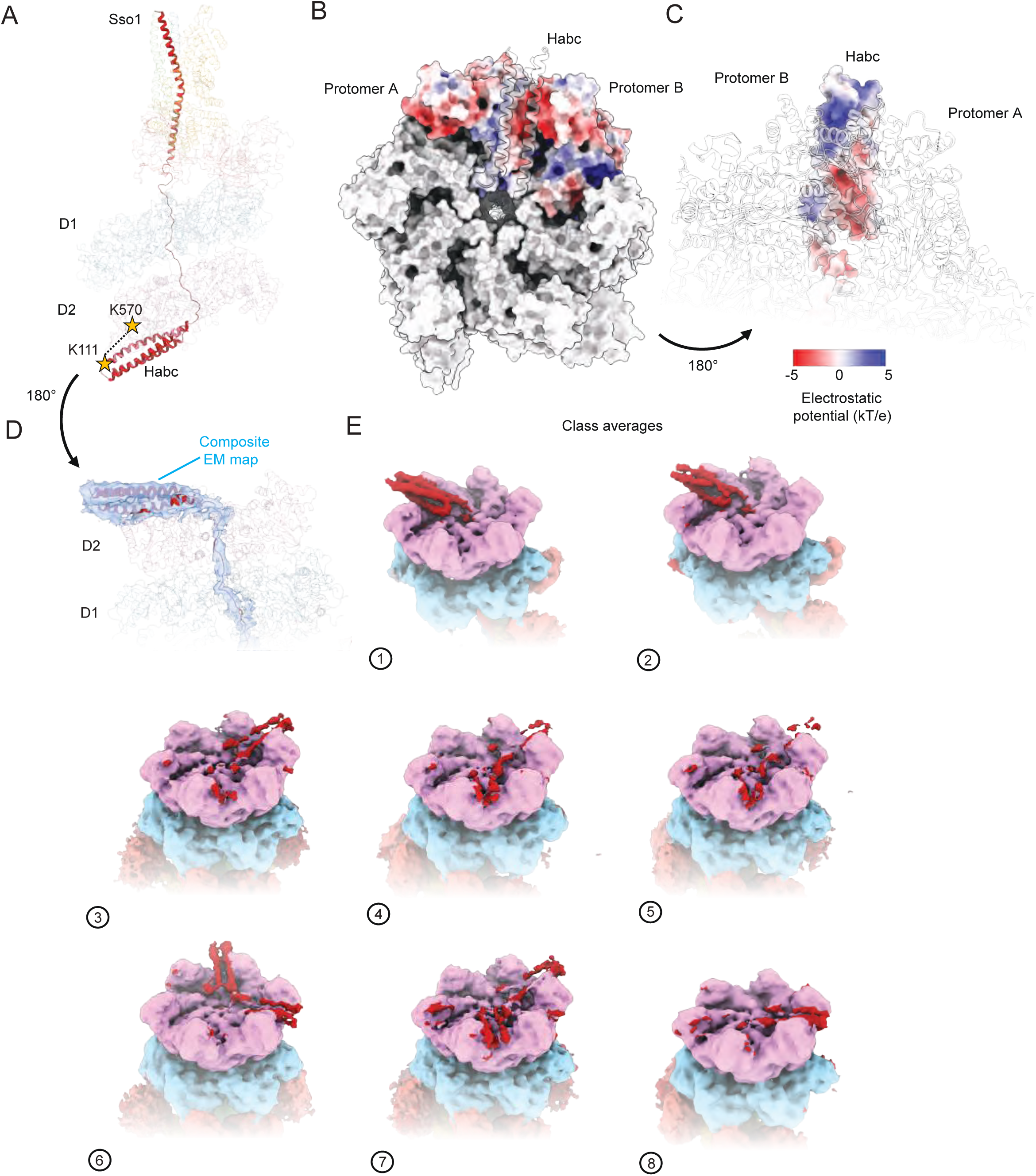
Sso1 is threaded through the D1 and D2 domains. **a,** y20S class 1 atomic model emphasizing the position of Sso1. Stars represent the crosslink identified by crosslinking mass spectrometry (Figure 1B). **b,** Bottom view of y20S class 1 D1 ring; the Habc domain is packed against the D2 ring surface, between protrusions corresponding to the C-terminal helical regions of two D2 small subdomains. All surfaces are colored grey, except for protomers A and B, which are colored by their electrostatic potential **c,** Rotated view of y20S class 1 relative to **b** showing the Habc tucked between protomers A and B with now the Habc domain colored by its electrostatic potential **d,** Composite cryo-EM map (blue, Relion-unsharpened map: Habc domain and D1 density and CryoSPARC 3DFlex map: D2 density) of y20S class 1. **e,** Cryo-EM maps of all eight y20S classes reveal the Habc domain associated with the Sec18 D2 ring in all classes (red). In 6/8 classes, the Habc density is radially averaged into multiple discrete positions around the D2 ring, leading to more diffuse density.

Remarkably, our cryo-EM structures reveal that Sso1 is loaded into both the D1 and the D2 pores since the Habc domain of Sso1 interacts with the outside of the D2 ring, with class 1 having the most well-defined, discrete density (**Figure 4A–E, Extended Data Figure 3B)**. In class 1, the density for Sso1 within Sec18 is continuous from the D1 pore entrance to the D2 pore exit, where the Habc domain begins (**Figure 4A, D**). The three-helix bundle of the Habc domain is then positioned outside the D2 pore, where it packs between two small D2 subdomains (protomers A and B) in an interaction driven by charge complementarity. Indeed, the electrostatic potential surface of the Habc domain forms a roughly opposite that of the bottom of the D2 surface against which it packs (**Figure 4B-C**). Given that this charge distribution is replicated between each pair of D2 subdomains around the D2 surface, it is unsurprising that the Habc domain adopts multiple corresponding rotational states over the eight classes, presumably due to the flexible linker that follows it (**Figure 4E**). This structural observation is consistent with XL-MS data showing a crosslink between the Habc and the D2 domain of Sec18 (**Figure 4A, Extended Data Figure 1B**). Although the putative distance between the crosslinked residues (K570 in Sec18 and K111 of Sso1) observed in the model for class 1 (42 Å) is at the limit for a crosslinking distance, the conformational flexibility of the Habc domain to rotate and exist in multiple conformations about the D2 ring (**Figure 4E**) likely produces conformations well within the crosslinking distance threshold.

### Sec18 does not unfold the Sso1/syntaxin Habc domain

Given that this complex was assembled in non-hydrolyzing conditions, we next asked how Sso1 is loaded into Sec18 given the diameter of the Habc domain of Sso1 is ∼30 Å, whereas the D1 pore of Sec18 has a diameter of ∼11 Å. In other AAA+ translocases like ClpX, ATP hydrolysis drives complete substrate threading through the AAA+ pore^27^. However, in the case of Sec18 in a non-hydrolyzing condition, such complete threading would require the Habc domain to unfold without any energetic input from Sec18.

We employed two orthogonal approaches to test if the Sso1 Habc domain unfolds during SNARE disassembly. In our first approach, we used a single-molecule Fluorescence Resonance Energy Transfer (smFRET) system developed previously for NSF, α-SNAP, and neuronal SNAREs^11^ in which individual SNARE domains were synthetically linked and stochastically labeled with FRET pairs. This system allows one to observe multiple rounds of SNARE disassembly as the disassembled SNARE domains readily reassemble into a *cis*-SNARE complex due to the covalent linkages. We chose this system since it is well-established. Moreover, it is relevant for the Sec18 system studied here since the primary sequence is conserved in the core regions of Sec18/NSF responsible for ATP hydrolysis and substrate processing (**Extended Data Figure 4A**). In addition, the D1 and D2 rings are structurally conserved when comparing the D1 and D2 rings of Sec18/NSF with engaged substrates, with an average Cα RMSD of 1.64 Å (**Extended Data Figure 4B-C**). The only protomer that exhibits much difference is the F protomer, likely due to its mobility and not due to inherent differences between the structures. Furthermore, we tested the ability of Sec18 and Sec17 to disassemble fluorophore-labeled neuronal SNARE and, *vice versa*, NSF and α-SNAP to disassemble fluorophore-labeled exocytic ySNARE complex (**Extended Data Figure 5A-E, Supplementary Discussion**). We found that both could process the species-ortholog SNARE complex, albeit at different rates, further corroborating the interchangeability of NSF and Sec18.

We thus used the smFRET assay to study the effect of NSF on the conformation of the three-helix bundle Habc domain of syntaxin during disassembly. We either stochastically labeled the SNARE domains or stochastically labeled the Habc domain at two distinct residue positions to monitor either SNARE disassembly or the folded state of the Habc domain (**Figure 5A**). As in our previous work^11^, we observed repeated rounds of disassembly and re-assembly for the SNARE domains as indicated by the changes in single FRET intensity over tens of seconds (**Figure 5B-C**). In stark contrast, we did not observe a change in single-molecule FRET intensity for the labeled Habc domain in non-disassembly and disassembly conditions, producing a FRET intensity distribution consisting only of a high-FRET state (the small peak at 0 is due to traces where acceptor photobleaching occurred since the particular traces never transitioned back to high FRET and no change in the peak intensity between non-disassembly and disassembly conditions was observed) (**Figure 5D-F**). This result suggests that the Habc domain does not unfold during NSF activity.

**Figure 5:**
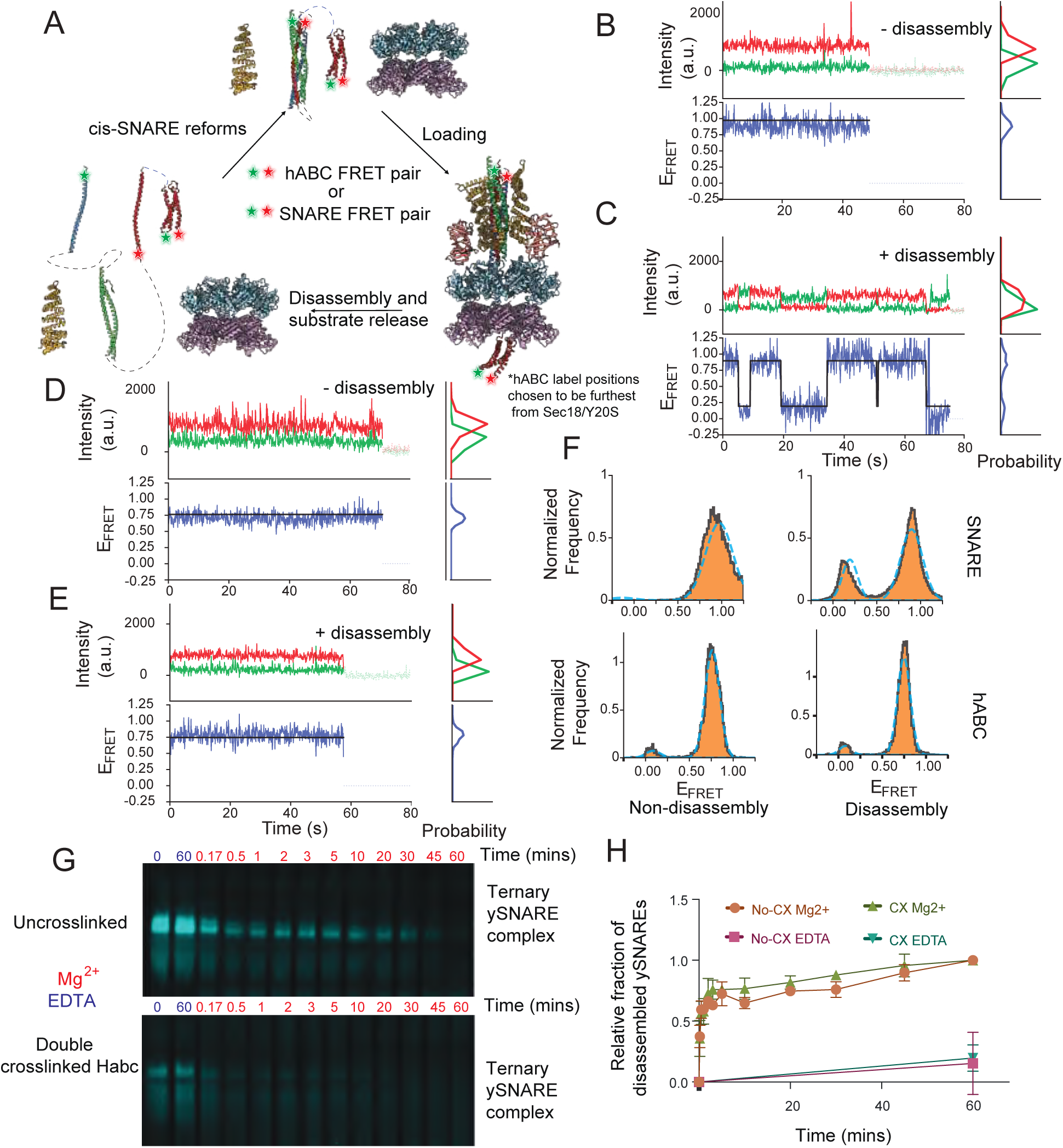
Sec18/NSF does not unfold the Habc domain. **a,** Diagram of the smFRET disassembly/re-assembly assay with NSF/α-SNAP and linked neuronal SNAREs. Either the linked SNAREs complex is stochastically labeled (residue number 249 in syntaxin and residue number 82 in synaptobrevin) or the syntaxin Habc domain is stochastically labeled (residue numbers 35 and 105 in the Habc domain). **b,** Representative traces of labeled linked SNARE complex in the non-hydrolyzing condition. Green represents the acceptor, and red represents the donor-dye fluorescence-intensity time traces. n = 53. **c,** Representative time trace of labeled SNARE complex in disassembly conditions, the black line represents fitting by a vbGMM^51^. n = 114. **d,** Representative time-trace of labeled Habc domain in the non-hydrolyzing condition. Green represents the donor, and red is the acceptor dye fluorescence intensity time-traces. n = 27. **e,** Representative time-trace of labeled Habc domain in disassembly conditions. The black line represents fitting a vbGMM. n = 22. **f,** Histograms of E-FRET for all four conditions. The blue dotted line represents a Gaussian fit of the data. The y-axis is the normalized frequency of occurrence of the E-FRET states in the traces. Top left: SNARE E-FRET for the non-hydrolyzing condition, top right: SNARE E-FRET for the hydrolyzing condition. Habc E-FRET for the non-hydrolyzing condition (bottom left). Habc E-FRET for the hydrolyzing condition (bottom right). **g,** Native gel of labeled uncrosslinked or crosslinked yeast SNARE complex that has been labeled with Oregon Green Maleimide 488. **h,** Quantification of gel densitometry of n = 3 independent disassembly experiments. Error bars represent the standard deviation of the mean. Densitometries were all normalized to 0 min values.

To corroborate this finding with an orthogonal approach for the yeast system, we double-crosslinked the Habc domain of Sso1 by incorporating an unnatural amino acid, 4-azido-L-phenylalanine^39^ (**Extended Data Figure 5F-G**). This double crosslink was introduced to prevent the Habc domain from unfolding during any Sec18-driven threading. After crosslinking, confirmed by mass-spectrometry in which we did not detect any uncrosslinked sample (**Extended Data Figure 5G**), we performed a disassembly assay on fluorescently labeled un-crosslinked and crosslinked complexes and monitored progress by native gel electrophoresis (**Figure 5G**). The top band, representing the fully assembled exocytic SNARE complex, is present in both conditions but disappears when adding Mg^2+^ to initiate hydrolysis and the disassembly reaction. There was no qualitative difference in the disassembly between uncrosslinked and crosslinked conditions after normalizing for labeling efficiency (**Figure 5H**). Thus, the folded state of the Habc domain does not change during yeast SNARE disassembly, and crosslinking the Habc domain does not affect the disassembly kinetics. These results for both the yeast and neuronal systems further argue against a threading model of Sso1 engagement since the Habc domain would be unable to pass through the D1 and D2 domains.

### Substrate-released Sec18 reveals coordinated ring opening

Together, these observations beg the question of how Sso1 enters the Sec18 ATPase rings. To address this topological challenge, we next determined structures of the y20S complex after initiating Sec18 hydrolysis by adding Mg^2+^ (referred to as “hydrolyzing condition”).

Informed by a fluorescent protein disassembly assay (**Extended Data Figure 5H)**, we initiated the disassembly reaction and waited 7 seconds before sample vitrification. In the resulting cryo-EM dataset, we found a small number of particles (6,356 or 2.26% of final particles, class 1) that consist of entire y20S assemblies (**Figure 6A**), largely similar to those observed in the non-hydrolyzing condition (**Figure 2)**. While the resolution of this reconstruction is low (10.88 Å), the D1 ring is flattened relative to reconstructions from the non-hydrolyzing condition (**Figure 6B**). Considering that our sample consists of purified y20S complexes before initiating disassembly by adding Mg^2+^, this D1 ring flattening likely occurs due to Mg^2+^ binding or ATP-hydrolysis at one or more subunits.

**Figure 6:**
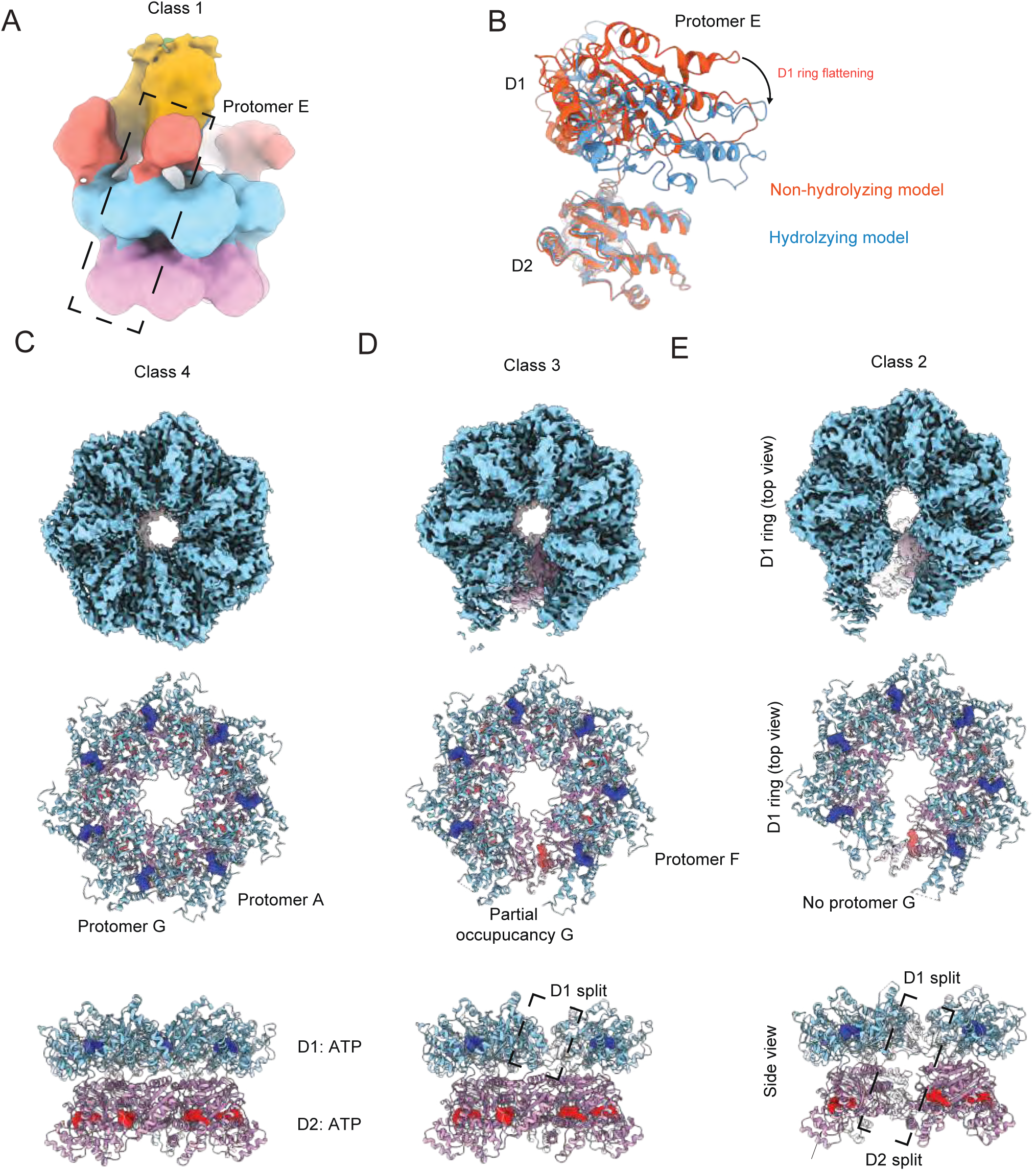
Post-disassembly of y20S reveals substrate-released states. **a,** Cryo-EM map of y20S in the hydrolyzing condition, colored by domain. **b**, The distance between the D1 and D2 domains decreases for some protomers under hydrolyzing conditions, leading to D1 ring flattening. The orange model corresponds to protomer E from the merged D1/D2 focused class in the non-hydrolyzing condition, which was rigid-body fit into the density of the hydrolyzing condition to generate the blue model. **c,** Cryo-EM map and model of class 4 of Sec18, showing a heptamer. **d,** Cryo-EM map and atomic model of class 3 of Sec18, suggesting a transition state. **e, Cryo-**EM map and atomic model of class 2 of Sec18, showing a split hexamer. In panels **c-e**, the top is the EM-map, while the middle and bottom show atomic models in top and side views, respectively.

The remaining particles (274,883 or 97.74 % of final particles) were substrate-free (classes 2–4 in **Figure 6C–E**). The D1 ring is also flat in all these substrate-free classes, with all D1 protomers bound to ADP. This uniform binding to ADP, as opposed to the spiral staircase of apo, ADP, and ATP-bound protomers, likely explains the flattened nature of the protomers. In the D2 ring, all protomers are still ATP-bound, consistent with their role in oligomerization. Class 4 consists of a configuration where both the D1 and D2 rings are heptameric (**Figure 6C**), and class 3 is similar to this heptameric class, except the 7th protomer is only partially occupied (**Figure 6D).**

Class 2 has a well-resolved hexameric configuration with a coordinated split in class 2 in the D1 and D2 rings (**Figure 6E**). This split spans ∼20 Å between protomers and is large enough to accommodate a polypeptide chain entering or leaving the rings. This conformational change is likely induced by the nucleotide state and specifically coupled to the presence of ADP throughout the ring following hydrolysis and SNARE substrate processing. The coordinated split in the D1 and D2 domain rings allows the processed SNARE substrate to be released from the side. This side-release mechanism thus addresses one part of the topological challenge stated above: threading occurs, but the substrate exits from the side, avoiding the membrane domains and linkages in SNARE proteins. We observed similar states, including a coordinate split of the D1 and D2 rings, for the mammalian homolog NSF after processing substrate^25^. This side-release mechanism may also explain the Snc2 crosslink to the residue in the D2 domain that is at the interface between the D1 and D2 protomers (**Figure 1F**).

### Substrate-free Sec18 reveals coordinated ring opening

Next, we asked if this same side-opening of Sec18 occurs in the hydrolyzing condition without SNARE substrate. We prepared purified wild-type Sec18 in the presence of ATP and Mg^2+^ in a three-stage procedure that yielded a pure sample used for cryo-EM studies (**Extended Data Figure 6A-C)**. Two bands were visible in native gels, indicating two oligomeric states (**Extended Data Figure 6D**). Processing and classification led to three classes, two of which appeared to be heptameric and identical (280,935 particles and 73,253 particles, respectively; **Figure 7A**) and the third class, which contained indeterminate density in the D1 ring (70,513 particles). Density for the N-domains was not well resolved in any of the classes, presumably due to the conformational flexibility of the N-domains without substrate or adaptor present.

**Figure 7:**
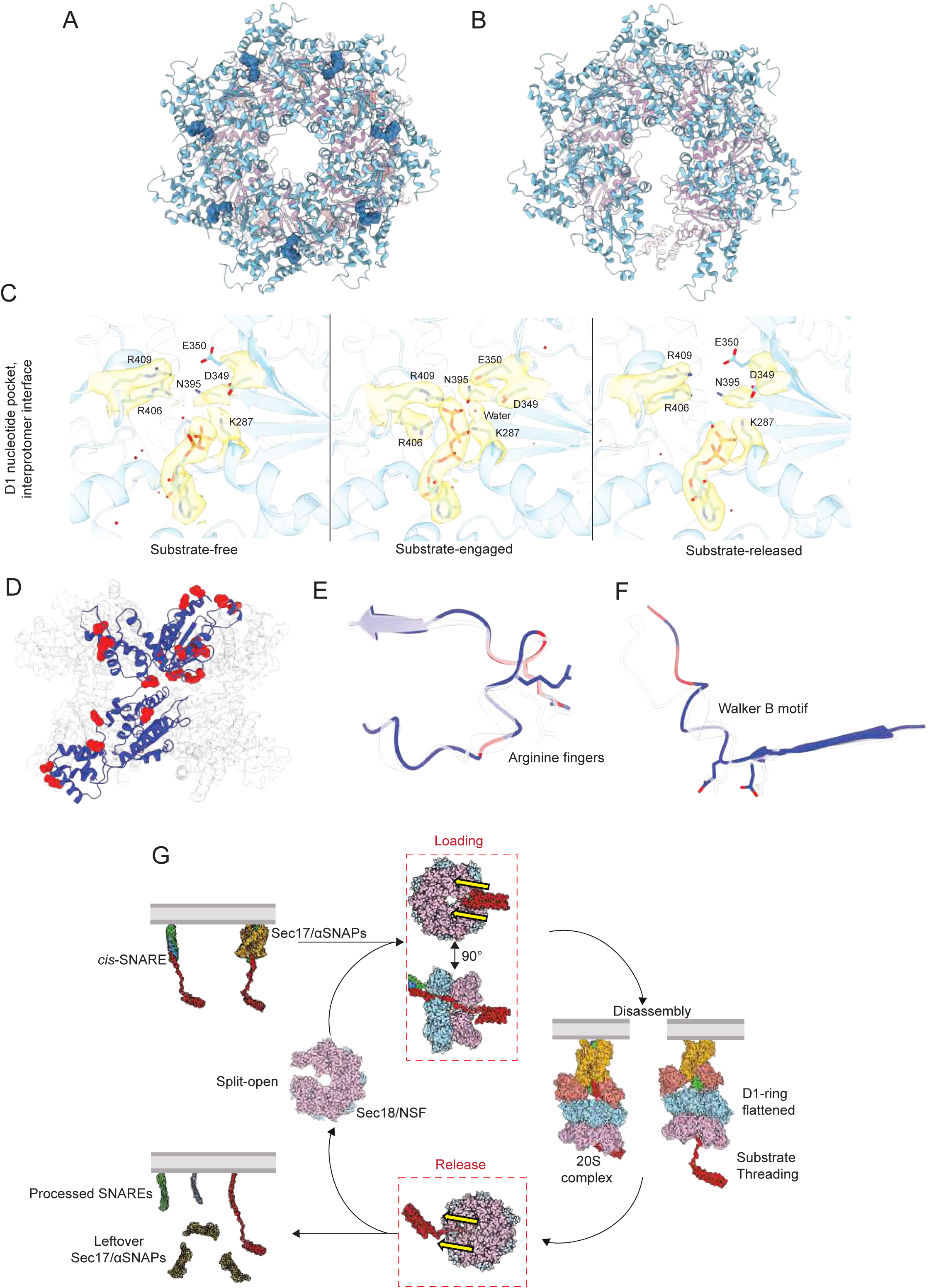
Structures of substrate-free Sec18 in the hydrolyzing condition. **a,** Atomic model of the substrate-free heptameric state of Sec18 (class 1). **b,** Atomic model of the split substrate-free hexameric state of Sec18 (derived from 3DVA of class 3). **c,** The D1 nucleotide binding pocket is remodeled as a function of SNARE substrate binding and nucleotide state. The arginine fingers R406 and R409 are contributed by the neighboring protomer. The substrate-free and substrate-released states are nearly identical in conformation. Water molecules are shown as red spheres. **d**, Atomic model of Sec18 with protomer A colored blue if not a significant residue from conformational analysis and colored red if significant (p-value < 0.05). **e,** Atomic models show the difference of the arginine finger loop in the no substrate (transparent) and substrate condition. **f,** Atomic models of the loop proximal to the Walker B motif in the no substrate (transparent) and substrate condition. For **d-f**, residues are colored red if they are in the top 5% of residues that significantly vary between conditions. **g.** We propose a general model of SNARE recycling, regardless of the cellular context. First, a *cis-*SNARE complex is coated with at least 1 Sec17/α-SNAP adaptor molecule, allowing Sec18/NSF to recognize it. The SNARE substrate is then loaded into a split hexamer with a coordinated opening in both D1 and D2 rings through the side, bypassing whatever N-terminal domains may be present. The substrate is then threaded coaxially through the Sec18/NSF pore. Upon completion of processing, the SNARE substrate is released through the side, bypassing the topological constraints of the membrane. Finally, Sec18 returns to its ‘resting’ state until more substrate is encountered. Yellow arrows indicate the direction of loading and release.

3D variability analysis (3DVA)^40^ of the third class suggests that it consists of a mixture of heptameric and hexameric Sec18 (**Extended Data Figure 6E),** consistent with the two bands seen on a native gel of the sample. The hexameric class reconstructed from 3DVA particle slices from the first mode (**Figure 7B**) is like class 2 observed for the y20S complex under hydrolyzing conditions (**Figure 6E**), wherein hexameric Sec18 has a large, coordinated split in its D1 and D2 domain rings without substrate. This similarity in Sec18 conformations provides further evidence that the substrate is both loaded and released through the side of the rings, explaining why the Habc domain is not unfolded during Sec18/NSF activity (**Figure 4**) and that crosslinking the Habc domain does substantially affect the disassembly kinetics (**Figure 5H**). This solves the second part of the topological question posed above.

The heptameric class (**Figure 7A**) is nearly identical to class 4 observed in the y20S complex under hydrolyzing conditions (**Figure 6C**). Each protomer in the heptamer is similar to the others, with an average RMSD of <1Å between them, with the angle between the D1 and D2 rings varying only with a small range of 61.9°–67.1°, in contrast to hexamer, with a range of 60.8°–92.3° (**Figure 2C, Extended Data Figure 6F**). The D1 pore diameter is 30 Å, larger than substrate-bound hexamer, with a diameter of ∼11 Å. The substrate-free structures (hexamer and heptamer) are nearly identical (RMSD = 0.48 Å) to the substrate-released structures.

To test if these D1 ADP-bound, ring-flattened states of Sec18 observed by single particle cryo-EM convert to a disassembly-competent state, we tested the same protein preparation used for cryo-EM with a vacuolar/lysosome proteoliposome fusion assay^41,42^ (**Extended Data Figure 6H-J, Supplementary Discussion**); this assay confirmed that the Sec18 preparation used for cryo-EM was active and processed vacuolar SNAREs. Moreover, NSF also adopts this flattened ring state and a heptameric or split hexameric arrangement when exchanged into a buffer containing ATP and Mg^2+^ without substrate (**Extended Data Figure 7, Supplementary Discussion**) and starting from neuronal 20S complex in a hydrolyzing condition^25^, consistent with our observations that Sec18 and NSF share the same functional mechanism.

### Implications for ATPase activity

The observed structures of Sec18 in substrate-free, substrate-loaded, and substrate-released states suggest large conformational change upon ATP hydrolysis, consistent with previous observations^24,25,43^. In the substrate-free and substrate-released states, examination of the D1 nucleotide binding pockets reveals critical catalytic residues far from the nucleotide (**Figure 7C).** However, in the substrate-engaged state, these conserved residues are tightly bound to nucleotide in a state preceding the binding of divalent cation required to drive hydrolysis. Employing the same statistical workflow used to cluster the different y20S classes (**Methods**), we identified the most significant residues that vary between the no substrate and substrate-bound state by generating an ensemble of models for each class via ensemble refinement (residues colored red in **Figure 7D**, variance contribution per component in **Extended Data Figure 5I**). For both the arginine finger loop and Walker B motif, we observed substantial changes to side chain and backbone conformations (**Figure 7E-F**), wherein coordinating residues are positioned closer to the nucleotide-binding pocket in the presence of substrate, presumably establishing a hydrolysis-ready state. In the case of the arginine fingers, the side chain conformational change is likely coupled to nucleotide identity, as hydrolysis disrupts ATP coordination, weakens the interprotomer interface, and leads to rigid body motion of the D1 large subdomain as the arginine fingers disengage. The case of the shift in the Walker B element, on the other hand, appears to be the flexing of the loop without substantial changes to the sidechains of D349 and E350, induced by the motions of pore loop 2 (GVG motif) that bind to the substrate in the pore.

### Mechanistic details of ring opening

The observed coordinated opening in both the D1 and D2 rings in conditions with and without substrate (**Figures 6 and 7**) suggests a side-release and side-loading mechanism of substrate. We speculate that the conformations of the split Sec18 observed in these multiple conditions are frequently sampled, enabling a substrate to enter or exit the D1-D2 double ring. To better understand the mechanism driving this opening, we compared the flat D1 closed and open conformations of Sec18 (**Figure 6C, E**). The Sec18 protomers are very similar (**Extended Data Figures 8A, 8B**), except for the protomers A and F at the split of the D1 and D2 rings. Rigid body motions alter the relative positions of these protomers’ respective sub-domains. We performed a conformational analysis to determine which amino acids contributed most to this conformational change between the open and closed states (**Methods**); the eight most significant residues contributing to the variance between the two conformations are shown in **Extended Data Figure 8C**. These residues are located in flexible regions between the structured sub-domains of a protomer (**Extended Data Figure 8D)**, where changes to the backbone ɸ and ψ angles of these residues lead to rigid body motions of subdomains. For example, four of the eight residues (R409, F410, L418, N690) are in the linker region between the large and small subdomains of the D1 domain, while G514 is situated between the D1 and D2 domains. These small and large subdomain movements, along with relative motion between the D1 and D2 domains, lead to the split that allows substrate loading or release. Given that this opening is hydrolysis independent and occurs regardless of the presence of substrate, it is likely that Sec18 samples these states upon thermal agitation; subsequently, substrate enters the pore and locks the D1 and D2 rings into an active hexameric form (*i.e.*, the 20S/y20S complex), possibly in concert with N-domain engagement.

## Discussion

Based on our LC-MS/MS, structural, and functional results, we propose the following model for SNARE recycling (**Figure 7G**), consisting of four phases. Sec18/NSF transiently forms split-open conformation of both D1 and D2 domain rings in the resting phase with no substrate. Once the *cis*-SNARE complex has been coated with adaptor Sec17/α-SNAP molecules, the Sec18 N-domains can bind the subcomplex and position the unstructured linker connecting the N-terminal Sso1 Habc domain to the SNARE domain proximal to the D1/D2 split without unfolding of the Sso1 Habc domain (**Figure 5H**), thus promoting the loading of substrate into the catalytic core of the enzyme. More generally, this side-loading mechanism would allow Sec18/NSF to process complexes incorporating all SNAREs in the cell regardless of the presence of folded N-terminal domains, provided that, in the latter case, a long enough linker connects them to the following SNARE domain. Upon forming the (y)20S complex, ATP-driven hydrolysis likely fuels the disassembly of the (y)SNARE complex by some degree of threading, accompanied by D1 ring flattening. In the case of Sso1, since full threading cannot occur due to the transmembrane domain, partial melting of the N-terminal end of the SNARE domain could be sufficient to destabilize the SNARE complex. Finally, the pore-bound SNARE is released through the side-split of both the D1 and D2 rings, regardless of membrane anchors or transmembrane domains in SNARE proteins. We observed a similar coordinated opening of the D1 and D2 rings for the mammalian homolog NSF after processing substrate^25^. The syntaxin Habc domain also does not unfold during NSF/mediated disassembly of the ternary SNARE complex (**Figure 5A-G**). The side-loading/release mechanism explains how Sec18/NSF can disassemble SNAREs with constrained topologies.

As observed in our cross-linking LC-MS/MS experiments (**Figure 1**), the Sec18 D2 ring can interact not just with the Habc domain of Sso1 but also with other N-terminal accessory domains (such as GOS1, TLG1, and TLG2). In turn, N-terminal domains of SNAREs often interact with other factors. For example, the trimeric Dsl1 complex binds to the SNAREs Use1, Ufe1, and Sec20 through interactions with N-terminal SNARE domains. This 255 kDa complex tethers endoplasmic reticulum membranes together before forming the *trans-*SNARE complex and likely remains complexed before, during, and after fusion^31^. Another example is HOPS, a hexameric complex that tethers lysosomal/vacuolar membranes and orchestrates their fusion^44^. HOPS has two Rab-binding subunits for tethering and a SNARE binding subunit that catalyzes initial SNARE complex assembly^45^. Sec18/NSF and Sec17/α-SNAP cooperate with HOPS to ensure rapid and efficient SNARE-mediated fusion^46,47^. For the case of Dsl1, the N-terminal domains of SNAREs remain in a supramolecular arrangement that must be bypassed. For vacuolar/lysosomal fusion, SNAREs must be preprocessed and handed off to the HOPS complex. These complexes impose topological constraints on both ends of the SNAREs that can be resolved by side-loading and side-release.

We observed heptameric states of Sec18 (**Figure 6C**) and NSF (**Extended Data Figure 7A**), split-open hexameric states of Sec18 (**Figure 6E**) and NSF^25^, and a transition between split-hexamer and heptamer of Sec18 (**Figure 6D**). Hydrolysis likely accelerates the transition from hexameric to heptameric state via the split-open state for two related reasons. First, while ring splitting itself is hydrolysis-independent, hydrolysis is associated with large-scale conformational change in the ATPase rings and likely increases the frequency of sampling the open state. Second, we have not observed the heptamer in a substrate-engaged state. Moreover, it seems unlikely that a heptamer could accommodate substrate given the changes heptamerization induces in the N- and D1-layers of the oligomer (an increased diameter would interfere with SNAP recognition of substrate and substrate engagement in the pore). As such, it seems reasonable that heptamer is enriched under hydrolyzing conditions either in the absence of SNARE substrate or in which a significant fraction of SNARE complexes have been disassembled—the hexamer samples the open state and the seventh protomer enters in the absence of SNARE substrate. Thus, the heptameric state, which is unable to process substrate, may not occur in the cell, as there is likely an abundance of *cis*-SNARE substrate and ATP to capture. Moreover, the XL-MS data cannot disentangle the presence of a hexamer or heptamer due to the small overall changes in residue distances between states. We conclude that NSF/Sec18 is mostly hexameric in the cell unless there is a significant dearth of substrate to process.

How broadly applicable is this AAA+ side-loading/release mechanism beyond NSF/Sec18? The NSF/Sec18 protein family is highly related to the p97/cdc48 family; both are clade 3 AAA+ machines with two AAA+ rings^13^. For p97/cdc48, their substrate is a polyubiquitinated molecule, again imposing topological restraints. In cryo-EM structures of cdc48 with substrate in non-hydrolyzing conditions, the substrate is threaded through the D1 and D2 rings, presumably without any ATP hydrolysis^48^. This study also showed that distal ubiquitin molecules on a substrate do not unfold during cdc48 activity. These observations suggest that this side-load and side-release mechanism may be employed by cdc48/p97 as well. Lastly, a split p97 structure was observed when bound to a cofactor^49^. This p97 structure also displayed a coordinated split in its D1 and D2 rings with flattened conformations, similar to those observed for Sec18, suggesting a conserved mechanism.

Finally, we note that distantly related AAA+ clamp loaders and their clamp also load their nucleic acid substrate from the side^50^. DNA is loaded into an open clamp and AAA+ clamp loader. Once the clamp loader recognizes the substrate, the ATPase is triggered, and the clamp loader closes the complex around the DNA. Once the clamp is fully secured, the clamp-loader is then released. This shared mechanism suggests the possibility of side-loading and release across the entire AAA+ protein family.

## Acknowledgments

We thank the late William I. Weis and Joseph D. Puglisi for stimulating discussions. We thank the National Institutes of Mental Health (NIMH) for support (RO1MH63105 to A.T.B., 1F31MH134477 to Y.A.K.). Y.A.K was additionally supported in part by an NSF GRFP and the Knight-Hennessy Scholarship. K.I.W. was supported by a postdoctoral fellowship from the Helen Hay Whitney Foundation, supported by the Howard Hughes Medical Institute. W.T.W. is supported by a grant from the NIH (2R35GM118037). We thank the Vincent Coates Foundation Mass Spectrometry Laboratory, Stanford University Mass Spectrometry (RRID:SCR_017801) for utilizing the Thermo Orbitrap Eclipse nanoLC/MS system (RRID:SCR_022212) that was purchased with funding from the National Institutes of Health Shared Instrumentation Grant 1S10OD030473, the Stanford Cancer Institute Proteomics/Mass Spectrometry Shared Resource (NIH P30 CA124435). This article is subject to HHMI’s Open Access to Publications policy. HHMI laboratory heads have previously granted a non-exclusive CC BY 4.0 license to the public and a sublicensable license to HHMI in their research articles. Pursuant to those licenses, the author-accepted manuscript of this article can be made freely available under a CC BY 4.0 license immediately upon publication.

## Author contributions

Y.A.K and A.T.B conceived and designed experiments. K.I.W, B.S., and E.M. assisted Y.A.K. with cryo-EM data collection. Y.A.K. performed and analyzed all experimental data. K.I.W. assisted Y.A.K. with cryo-EM data processing and model building. R.A.F. and L.E. assisted with protein purification. G.M., F.L., and T.M. assisted with high-resolution MS proteomics data collection and analysis. K.D. assisted with bulk disassembly assays. U.B.C. collected and assisted with smFRET data. W.T.W. performed and analyzed vacuolar fusion assays. Y.A.K., A.T.B., and K.I.W. wrote the manuscript.

## Methods

### HA-Sec18 *S. cerevisiae* strain preparation and treatment

Briefly, we created an *S. cerevisiae* strain containing HA-tagged Sec18 and treated it with zymolyase and lyticase to dissolve its cell wall but maintain its integrity as a living cell (spheroplast). We then incubated the spheroplasts with disuccinimidyl glutarate (DSG), a membrane permeable protein crosslinking agent, for 30 minutes before cellular lysis and anti-HA co-immunoprecipitation (coIP) to isolate Sec18 and any bound proteins to it. This sample was then subjected to SDS-PAGE gel electrophoresis, and the bands corresponding to crosslinked Sec18 were excised and sent for liquid chromatography-mass spectrometry (LC-MS/MS) analysis. More specifically, tagged Sec18 (pYK175, 3x-Flag N-terminal, HA-tag C-terminal) plasmid was transformed into *S. cerevisiae* S288C using the Frozen-EZ Yeast Transformation Kit (Zymo Research T2001). Expression of tagged Sec18 was accomplished by growing yeast in 1% raffinose and 1% galactose media. Cells were spun down and treated with 100 units of zymolyase (Zymo Research #1004) at 37°C for 1 hour. After incubation, spheroplasts were pelleted with a 1000 xg spin. DSG diluted in 1x PBS was used to resuspend the pellet, and the sample was incubated for 30 minutes. The excess DSG was then quenched with 1M Tris and incubated for another 15 minutes. Cells were washed with cold PBS and were resuspended in lysis buffer (5mM EDTA, 1mM ATP, 1mM TCEP, 150 mM NaCl, 10% glycerol, 0.5% NP40, and 50 mM HEPES pH 7.4) and glass beads (BioSpec 11079105). This bead-lysis-cell solution was vortexed for 5 minutes at 4°C three times with ice incubations between each vortexing. The solution was then spun down at 10,000 xg for 5 minutes at 4°C. The supernatant was taken and then spun down again at 10,000 xg for 15 minutes at 4°C. The supernatant was then taken and subjected to magnetic-bead anti-HA coIP (Thermo Scientific 88838). The eluant of the beads was run on an SDS-PAGE gel, and the bands corresponding to Sec18 were excised.

### XL-MS sample preparation

Protein samples were embedded in Coomassie-stained gel bands, fixed in 1% acetic acid. The gels were washed several times in 50 mM ammonium bicarbonate, followed by reduction with 10 mM of dithriothreitol (DTT) and incubation for twenty minutes at room temperature. They were then alkylated using 30 mM acrylamide for 30 minutes. This was proceeded by overnight digestion at 37 degrees Celsius with 500ng of mass spectrometry grade trypsin/LysC mix (Promega). Post-digestion, samples were quenched with formic acid (adjusted to a pH ∼3) and desalted using MonoSpin C18 Solid-Phase Extraction (SPE) columns (GL Sciences). Finally, the samples were dried via SpeedVac (Thermo Scientific, San Jose, CA) and exchanged into LC-MS reconstitution buffer (2% acetonitrile with 0.1% formic acid in water) for instrumental analysis.

### LC-MS/MS analysis

Mass spectrometry experiments were performed using an Orbitrap Eclipse Tribrid mass spectrometer (Thermo Scientific, San Jose, CA) attached to an Acquity M-Class UPLC system (Waters Corporation, Milford, MA). The UPLC system was set to a flow rate of 300 nL/min, where mobile phase A was 0.2% formic acid in water and mobile phase B was 0.2% formic acid in acetonitrile. The analytical column was prepared in-house with an I.D. of 100 microns pulled to a nanospray emitter using a P2000 laser puller (Sutter Instrument, Novato, CA). The column was packed with Dr. Maisch 1.9 micron C18 stationary phase to a length of approximately 25 cm. Peptides were directly injected into the column with a gradient of 3-45% mobile phase B, followed by a high-B wash over a total of 80 minutes. The mass spectrometer was operated in a data-dependent mode using CID fragmentation for MS/MS spectra generation.

The RAW data were analyzed using Byonic v5.2.5 (Protein Metrics, Cupertino, CA) to identify peptides and infer proteins. Initial Byonic analyses used a concatenated FASTA file containing the Uniprot *Saccharomyces cerevisiae* proteins and other likely contaminants and impurities. Once sample complexity was determined, a second round of Byonic analyses was completed using a targeted FASTA file, which included only the sequences present in the samples and allowances for crosslinked peptides with the appropriate linker. Proteolysis with Trypsin/LysC was assumed to be fully specific with up to two missed cleavage sites. The precursor ion tolerance was set to 12 ppm. The fragment ion tolerance was set to 0.4 Da. Cysteine modified with propionamide was set as a fixed modification in the search. Variable modifications included oxidation on methionine and acetylation on protein N-terminus. Proteins were held to a false discovery rate of 1% using the standard reverse-decoy technique.

The following procedure validated Potential crosslinked peptides using Byologic v5.2.31 (Protein Metrics). Given the number of possible crosslinked peptides observed in these experiments, additional empirical constraints were applied to the potential crosslinked peptides to produce a more rigorous validation set for comparison with other biochemical assays. For the BSG cross-linking studies, crosslinked spectra were required to meet the following criteria: 1) all peptides, crosslinked or native, were filtered to a <1% FDR; 2) precursor mass error of less than 7 ppm was required for crosslinked peptides, and 3) the peptide primary sequence was at least 6 amino acids in length for at least one of the crosslinked peptide pairs. For zero-length crosslinking data analysis, the following additional constraints were added to those described above for BS3 crosslinking: 1) a minimum length of five amino acids was required for both members of the crosslink; 2) an alternative ‘XLink’ algorithm from Byonic was used to make assignments based on fragmentation of both peptides rather than just crosslink partner mass; 3) the crosslinks were assumed valid only if the protein contained the lysine crosslink partner; and 4) at least two crosslinked peptide spectra were assigned to the linkage. Following this, identified crosslinked peptides were further categorized into three groups: high confidence, medium confidence, and low confidence, where “high confidence” succeeded on all these rules and “low confidence” failed at least in three of these rules.

### Preparation of y20S in the non-hydrolyzing condition

The y20S complex consists of the Sso1–Snc1–Sec9 SNARE (ySNARE) complex, the Sec18 oligomeric complex, and Sec17 adaptor proteins, and it was assembled in a multi-stage workflow. Starting with the yeast exocytic complex, individual SNARE components Sec9 (401-651), Sso1 (1-265), and Snc1 (1-93) were purified individually. All three proteins were cloned into the pET28b plasmid vector with a 6x-His tag and a TEV cleavage sequence N-terminal to the SNARE protein sequence.

For Sec9, we used a lysis buffer of pH 7.5, 50 mM NaPi, 300 mM NaCl, 10 mM Imidazole, 0.5 mM TCEP, 1% Triton X-100 and SIGMAFAST Protease Inhibitor Cocktail Tablet (Sigma-Aldrich S8830) which was used to resuspend bacterial cell pellets from an 8L culture of auto-induced One Shot BL21 (DE3) *E*. coli (Thermofisher C600003). This resuspended lysate was subjected to sonication for 20 minutes (3 seconds on, 9 seconds off, 60% amplitude), followed by a 30-minute 4000 xg spin and a 1-hour 40,000 xg spin. The supernatant was bound (1 hour, 4°C), run through a Ni-NTA column, and washed with pH 7.5, 50 mM NaPi, 30 mM Imidazole, 300 mM NaCl, and 0.5 mM TCEP buffer. The sample is then eluted with a pH 8 50 mM NaPi, 400 mM Imidazole, 300 mM NaCl, and 0.5 mM TCEP buffer. This sample is then digested overnight with TEV protease in a dialysis buffer (pH 8 20 mM Tris, 250 mM NaCl, 0.5 mM TCEP, and 1 mM EDTA). The cleaved sample was then subjected to a MonoS 5/50 GL with buffer A consisting of pH 8 20 mM Tris, 50 mM NaCl, 0.5 mM TCEP, 1mM EDTA and with buffer B consisting of pH 8 20 mM Tris, 500 mM NaCl, 0.5 mM TCEP, and 1mM EDTA. Taking the majority peak corresponding to Sec9, we then subjected this peak to size exclusion chromatography (SEC) with the HiLoad 16/60 Superdex 200. The peak corresponding to Sec9 was concentrated and flash-frozen. We followed the same protocol for individually purifying Snc1 and Sso1.

Once all three individual SNAREs were purified, a 1:1:1 molar ratio of these proteins was added to a 6M GdHCl solution. This solution was then slowly dialyzed overnight at 4°C into a solution of 250 mM NaCl, 50 mM HEPES pH 7.6. This sample was then diluted to a NaCl concentration of 75 mM and then subjected to MonoQ with the low salt buffer at 75 mM and the high salt buffer at 500 mM with both buffers in 50 mM HEPES pH 7.6. The peak corresponding to a 1:1:1 ratio of all three bands, indicative of the yeast exocytic complex, was then concentrated for y20S formation without freezing.

The complete Sec18 protein coding sequence was cloned into the pMZ0002/pYK103 backbone with a 6x-His tag and TEV cleavage sequence N-terminal to Sec18. The 8L culture of One Shot BL21 (DE3) *E.* coli (Thermofisher C600003) was spun down and resuspended in lysis buffer of pH 7.5 100 mM HEPES, 500 mM KCl, 5 mM ATP, 5mM MgCl_2_, protease inhibitor tablets, and benzonase. This lysate was sonicated for 20 minutes (3 seconds on, 9 seconds off, 60% amplitude) and clarified with a 45,000 xg spin for 30 minutes. The lysate was supplemented with 20 mM Imidazole bound to Ni-NTA beads (4°C) and washed with pH 7.5 20 mM HEPES, 480 mM KCl, 0.5 mM ATP, 0.5 mM TCEP, 20 mM Imidazole, 1 mM MgCl_2_, and 10% glycerol. The sample was then eluted with pH 7.5 20 mM HEPES, 480 KCl, 0.5 mM ATP, 0.5 mM TCEP, 500 mM Imidazole, 1 mM MgCl_2,_ and 10% glycerol. The protein-containing peak was then equilibrated in SEC buffer pH 6.8 20 mM PIPES, 125 mM KCl, 0.2M sorbitol, 5 mM MgCl_2_, 2 mM ATP, 2 mM DTT, and 10% glycerol and injected a HiLoad 16/60 Superdex 200. The fractions were pooled, concentrated, and snap-frozen for later usage. The band right below Sec18 is an oligomeric *E. coli* contaminant, which is removed in the final steps of y20S complex purification (**Figure S2A, C**). Moreover, this contaminant is absent in the protocol for preparing high-purity Sec18 (see below and **Figure S6B**).

The full Sec17 coding sequence was cloned with a 10x-His tag and a TEV cleavage sequence fused N-terminally. The 8L culture of One Shot BL21 (DE3) *E. coli* was spun down and resuspended in a lysis buffer of pH 8 50 mM NaPi, 300 mM NaCl, 20 mM Imidazole, and 0.5 mM TCEP. This mixture was then sonicated (3 seconds on, 9 seconds off, 60% amplitude) and clarified at 45,000 xg for 30 minutes. This supernatant was then incubated with Ni-NTA beads for an hour and then washed with lysis buffer. The protein was eluted with lysis buffer supplemented with 500 mM Imidazole and 5 mM EDTA. TEV protease was added to pooled fractions and slowly dialyzed overnight at 4°C against a pH 8 50 mM Tris, 100 mM NaCl, 1 mM EDTA, 0.5 mM TCEP buffer. This sample was then diluted to a NaCl concentration of 50 mM and run on a MonoQ 10/100 from a 50 mM to 500 mM salt gradient. The major peak corresponding to Sec17 was then pooled and injected into a HiLoad 16/60 Superdex 200 SEC. The major peak from this run was taken, pooled, and snap-frozen for later usage.

To assemble the entire y20S complex, Sec17 was added to the freshly prepared yeast exocytic SNARE complex, followed by EDTA quenched Sec18 with a final ratio of Sec18:SNARE:Sec17 at 1:1.67:10, where a total of 5580 picomoles of Sec18 was used, in the following buffer: pH 8 50mM Tris, 150 mM NaCl, 1 mM TCEP, 1 mM ATP, and 1 mM EDTA. Following this assembly, this mixture was injected onto a Superose 6 10/300 increase column, and the peak corresponding to the y20S complex, as assessed by elution volume and corresponding SDS-PAGE gel, was concentrated to ∼40 mg/mL.

### Single particle Cryo-EM grid preparation of y20S in the non-hydrolyzing condition

Quantifoil R1.2/1.3 200 mesh gold grids were treated with chloroform and dried overnight. Grids were glow discharged, and 5 microliters of the sample (at a final concentration of 20 mg/mL with 0.05% v/v Nonidet P-40) was blotted onto the grids and further blotted and vitrified in liquid ethane using an FEI Vitrobot (ThermoFisher Scientific). Grids were blotted for 4 seconds with a 5-second wait time (reduced to 3 seconds in hydrolyzing condition).

### Sample preparation and single particle Cryo-EM of y20S in the hydrolyzing condition

Samples were prepared identically to y20S in the non-hydrolyzing condition, except that excess MgCl_2_ was blotted onto grids immediately before vitrification, and the total blot and wait time was reduced to 7 seconds before freezing.

### Sample preparation and single particle Cryo-EM of substrate-free Sec18 in the hydrolyzing condition

An alternative protocol was required to prepare high-purity, substrate-free Sec18 since low-purity Sec18, which can form y20S, was not amenable to high-quality cryo-EM studies. Therefore, we used a different protocol for preparing NSF^10^, except that at the final re-assembly step, we used MgCl_2_ instead of EDTA. The cryo-EM studies were performed similarly to the studies of Y20S in the non-hydrolyzing condition.

### Sample preparation and single particle Cryo-EM of NSF in the hydrolyzing condition

A sample of high-purity NSF was prepared, as described in the previous section. This sample was exchanged into a buffer supplemented with 1mM magnesium chloride instead of EDTA and incubated before a final SEC and freezing.

### Single particle cryo-EM data collection, processing, and model building

Single particle cryo-EM data collection and processing workflow are described thoroughly in the **Extended Table 1** and **Supplementary Figure 2**. An FEI Titan Krios (ThermoFisher Scientific) cryo-electron microscope with either a K3 Gatan or a Falcon F4i camera was used for data collection. Micrographs were analyzed using a combination of initial Relion 3.1^52^ processing followed by additional rounds of 3D classification or heterogenous refinement, and final refinements in CryoSPARC v4^53^. A general workflow description is provided in **Supplementary Figure 2**, and specific dataset information is provided in **Supplementary Figures 2-6** as well.

y20S models were constructed by first docking the crystal structure of the exocytic SNARE complex (PDB ID: 3B5N) and PDF/AlphaFold2^54^ models for Sec17 (1QQE/AF-P32602-F1-v4) into the unsharpened density first obtained in Relion 3.1^52^. Next, sharpened cryoSPARC-v4^53^ maps were used to model the Sec18 complex and the region of Sso1 that extended from the spire into the D1 pore. For class 1, 3DFlex analysis in cryoSPARC-v4 was performed. A mixture of the Relion 3.1 and cryoSPARC-v4 unsharpened and sharpened maps, and the 3Dflex map were used to model the domain outside of the D2 ring. The resulting atomic models were iteratively refined with a combination of COOT^55^, Phenix real-space refinement^56^, and ISOLDE^57^.

Substrate-free Sec18 and NSF models were built *de novo* in COOT using cryoSPARC sharpened maps and iteratively refined with ISOLDE and Phenix. AlphaFold2 model of Sec18 was used as a starting reference for model building (AF-P18759-F1-v4).

### Structural and conformational analysis

Statistical structural analysis was performed to identify the most significant backbone motions between the different structural states of Sec18 without substrate, as observed in Figure 6. Using Python 3, principal component analysis (PCA) on backbone ɸ/ψ angles was performed, and the single mode that contributed most to the variance (>70%) was analyzed. y20S models were then clustered in this reduced 5-dimensional space. When comparing the substrate-free and substrate-engaged states, ensembles consistent with a given map were generated to avoid model-building bias using Phenix’s simple molecular dynamics implementation for 2000 steps followed by a round of phenix.refine^56^. These ensembles were then subjected to backbone ɸ/ψ PCA, and the mode that contributed to the largest variance between the two clusters for the two classes was used to determine which residues contributed most to the difference in structures. For more details, please see the conformational tool package deposited in GitHub for annotated and commented code (https://github.com/YousufAKhan/ConformationalAnalysis/).

### smFRET disassembly/re-assembly assay

smFRET assays with linked neuronal SNAREs, NSF, α-SNAP, and reagents were prepared, as previously reported^11^. Briefly, linked SNARE complex (L-SNARE) composed of SNAP25A, synaptobrevin-2, and syntaxin-1A were biotinylated and flowed onto the chamber that was passivated by 10 mg/mL egg phosphatidylcholine 50 nm liposomes coated with 1 mg/mL of biotinylated BSA to mimic the lipid environment inside the cell. 0.1 mg/mL streptavidin was added to surface-tether the biotinylated and labeled L-SNARE complexes. L-20S was then assembled by first adding αSNAP and then later NSF (if in disassembly conditions). L-SNARE was diluted such that about 500 molecules of L-SNARE per 45 x 90 µm^2^ field of view were visible. The labels for the Habc domain are at residue positions 35 and 105, and labels for the L-SNARE complex are at residue position 249 of syntaxin and residue position 82 of synaptobrevin, identical to a previous study^11^. The Habc label positions are at the tips of the α-helices that are positioned away from the D2 pore and the Sec18 complex (Fig. 5a). Thus, one would expect that 20S complex formation does not affect the quantum yield and conformational dynamics of the dyes, which is indeed what we observe. The labeling was performed by mutating the residues at these positions to cysteines and stochastic conjugation with Alexa 555 and Alexa 647 dyes.

All smFRET experiments were performed on a prism-type total internal reflection fluorescence (TIRF) microscope using 532 nm (green) laser (CrystaLaser) excitation. Two observation channels were created by a 640 nm single-edge dichroic beamsplitter (FF640-FDi01-25×36, Shemrock): one channel was used for the fluorescence emission intensity of the Alexa 555 dye and the other channel for the Alexa 647 dye. The two channels were recorded on two adjacent rectangular areas (45 × 90 μm2) of a charge-coupled device (CCD) camera (iXon+ DV 897E, Andor Technology USA, South Windsor CT). The imaging data were recorded with the smCamera program (Taekjip Ha, Johns Hopkins University, Baltimore). Flow chambers were assembled by creating a “sandwich” consisting of a quartz slide and a glass coverslip that were both coated with polyethylene glycol (PEG) molecules consisting of 0.1% (w/v) biotinylated-PEG except when stated otherwise and using double-sided tape to create up to five flow chambers.

Fluorescence intensity time traces were recorded by the smCamera^58^ program, peaks were selected, and correlated in the two channels as previously described^19^. The time traces were imported into tMaven^51^ for smFRET analysis. The standard, recommended tMaven workflow was employed. In short, all traces were evaluated manually and not subject to any automated trace preprocessing in tMaven. Traces that did not have single photobleaching events were not considered. After obtaining all traces that fit these criteria, the automatic photobleaching algorithm was used to remove regions that were photobleached for analysis of the FRET states.

Following this, the vbGMM model was used to model the transition between states with an assumption of two states (i.e., high and low). Following this, the tMaven “one dimension histogram” tool was used to generate histograms of the distribution of states of the raw FRET efficiency E=I_A_ / (I_A_+I_D_), where I_A_ and I_D_ are the acceptor and donor fluorescence intensities, respectively; no corrections for bleed-through, detection efficiency, and quantum yield were applied.

### Crosslinking assays

Cysteines were introduced into Sec9 and Sso1 for labeling with Oregon Green Maleimide 488 (O6034). UAG stop codons were introduced at V57 and F83 in Sso1 for crosslinking. Cys-Sec9, Cys-Sso1 (with or without stop codon depending on if used for crosslinked), and 6x-His-TEV-Snc1 were co-expressed (if crosslinked, also transformed with pEVOL-pAzF AddGene #31186, 2mM AzF and 0.2% v/v arabinose) and induced with IPTG. Cell pellets were lysed with pH 7 50 mM Tris, 200 mM NaCl, 1% v/v Triton X-100, 20 mM Imidazole, 0.5 mM TCEP and protease inhibitor tablets. Lysate was sonicated for 20 minutes (3 seconds on, 9 seconds off, 60% amplitude) and then clarified at 40,000 xg for 30 minutes. Following a 1-hour incubation at 4°C, the beads were washed with two different buffers. Buffer 1 is simply a lysis buffer without Triton X-100, and Buffer 2 is a high salt wash of pH 7 50 mM Tris, 1M NaCl, 50 mM Imidazole, 0.5 mM TCEP. The complex was eluted with pH 7 50 mM Tris, 200 mM NaCl, 350 mM Imidazole, and 0.5 mM TCEP. The sample was then digested overnight with TEV protease and dialyzed into pH 7 20 mM Tris, 200 mM NaCl, 0.5 mM TCEP. This sample was then injected onto a HiLoad 16/60 Superdex 200. The fractions containing all three in equimolar ratios were taken and concentrated. Samples were labeled with Oregon Green Maleimide 488 in molar excess overnight before cleanup with a Zeba spin column (ThermoFisher 89889). If the sample was meant to be crosslinked, it was subjected to UV irradiation before Oregon Green Maleimide labeling. The buffers used for these crosslinking assays were the same as the final dialysis buffer but were degassed before UV irradiation and Oregon Green Maleimide labeling.

For native gel disassembly assays, reactions were assembled in test tubes, initiated with MgCl_2_, and quenched with EDTA. Samples were then loaded into Any kD Mini-Protean gels (Bio-Rad 4569033) and used Tris-glycine running buffer (Bio-Rad) supplemented with 1 mM ATP to prevent Sec18 from falling apart in the gel. Gels were first visualized in an iBright 1500 (ThermoFisher Scientific) to visualize ySNARE bands only before coomassie staining. Gel densitometry was assessed in ImageJ.

### FRET and fluorescent dequenching disassembly assays

Neuronal disassembly assays were carried out as previously reported^10^, leveraging fluorescence dequenching of the neuronal SNARE complex labeled with Oregon Green Maleimide 488. For yeast SNARE disassembly assays, a strategy was devised to tag Snc1 with mTurq and Sso1 with mVenus. When assembled, excitation by a 434 nm laser leads to mVenus emission. 434 nm laser excitation leads to mTurq emission. Disassembly is followed by monitoring excitation/emission at 434/474 nm such that as the SNARE complex is disassembled, mTurq emission increases. We used the same instrument (FlexStation II 384, Molecular Devices) and a 384-well format to perform both the dequenching and FRET assays that were initiated by titrating MgCl_2_ in excess of the reactions.

### Vacuolar fusion assay

Assays were performed as previously reported^41,42^ with the same substrate-free Sec18 protein sample/batch that was purified and used for Cryo-EM. Briefly, proteoliposomes with vacuolar SNAREs were prepared with either Phycoerythirin-biotin (PhycoE) or Cy5 and incubated in a fluorescence plate reader for 40 minutes. FRET between PhycoE and Cy5 was measured at 1-minute intervals. Reaction mixtures contained HOPS complex, Sec17, and variable amounts of Sec18.

### Data availability statement

The mass spectrometry proteomics data have been deposited to the ProteomeXchange Consortium via the PRIDE partner repository (https://www.ebi.ac.uk/pride/) with the dataset identifier PXD062002 and 10.6019/PXD062002. All cryo-EM maps and coordinates have been deposited in the EMDB and PDB, respectively. The codes for the following classes (PDB/EMD)

- Y20S non-hydrolyzing condition

`o` Class 1

▪ 9CRU/EMD-45883
`o` Class 2

▪ 9N22/EMD-48826
`o` Class 3

▪ 9CRX/EMD-45885
`o` Class 4

▪ 9NG2/EMD-49380
`o` Class 5

▪ 9NLU/EMD-49522
`o` Class 6

▪ 9NLW/EMD-49524
`o` Class 7

▪ 9NLY/EMD-49526
`o` Class 8

▪ 9NLZ/EMD-49527
`o` Class 9 (All merge, focus on D1/D2)

▪ 9NM1/EMD-49528
- Y20S hydrolyzing condition

`o` Class 1

▪ No model created/EMD-70472
`o` Class 2

▪ 9NUD/EMD-49801
`o` Class 3

▪ 9NUE/EMD-49802
`o` Class 4

▪ 9NUZ/EMD-49824
- Sec18 hydrolyzing condition

`o` Class 1

▪ 9NV1/EMD-49826
`o` Class 2

▪ 9NV0/EMD-49825
- NSF hydrolyzing condition

`o` Class 1

▪ 9NV9/EMD-49831
`o` Class 2

▪ 9NVD/EMD-49833

The smFRET data have been deposited in the Stanford Data Repository (https://purl.stanford.edu/bm031pm6709). All data are available with the manuscript online and materials can be obtained from AddGene.org, where all the constructs will be deposited.

### Software and code availability

The conformational tool package has been deposited in GitHub at https://github.com/YousufAKhan/ConformationalAnalysis and smFRET related scripts can be found at https://github.com/brungerlab/single_molecule_matlab_scripts.

---

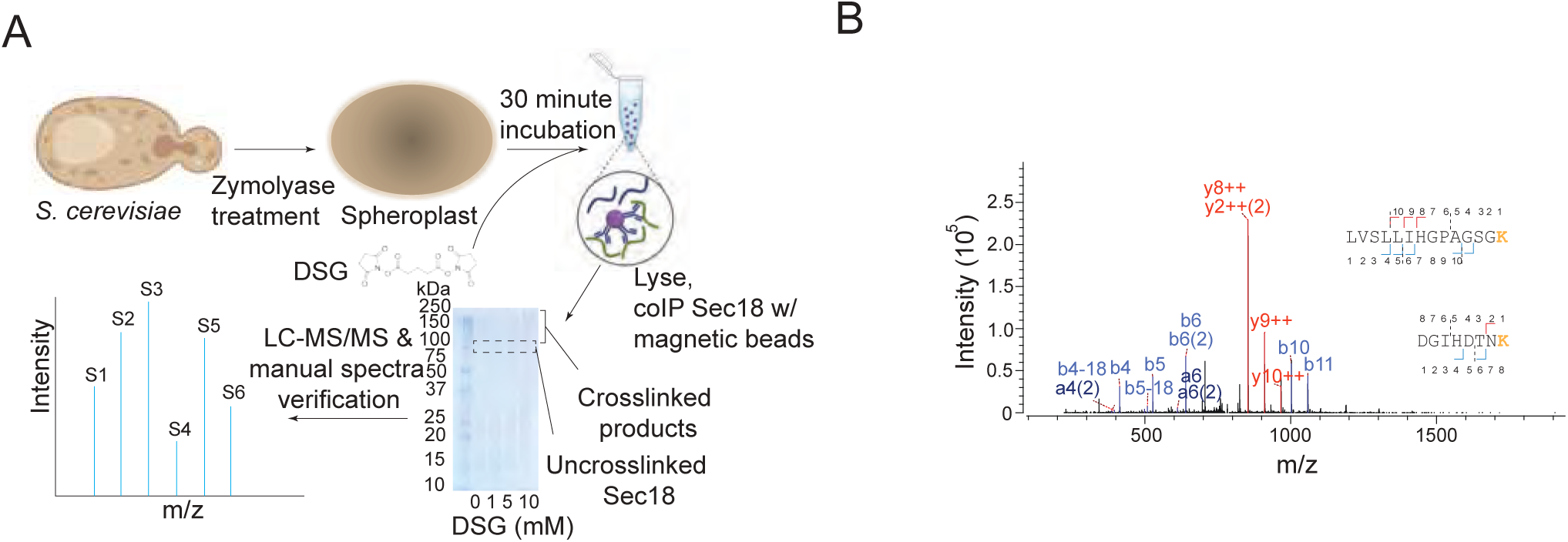

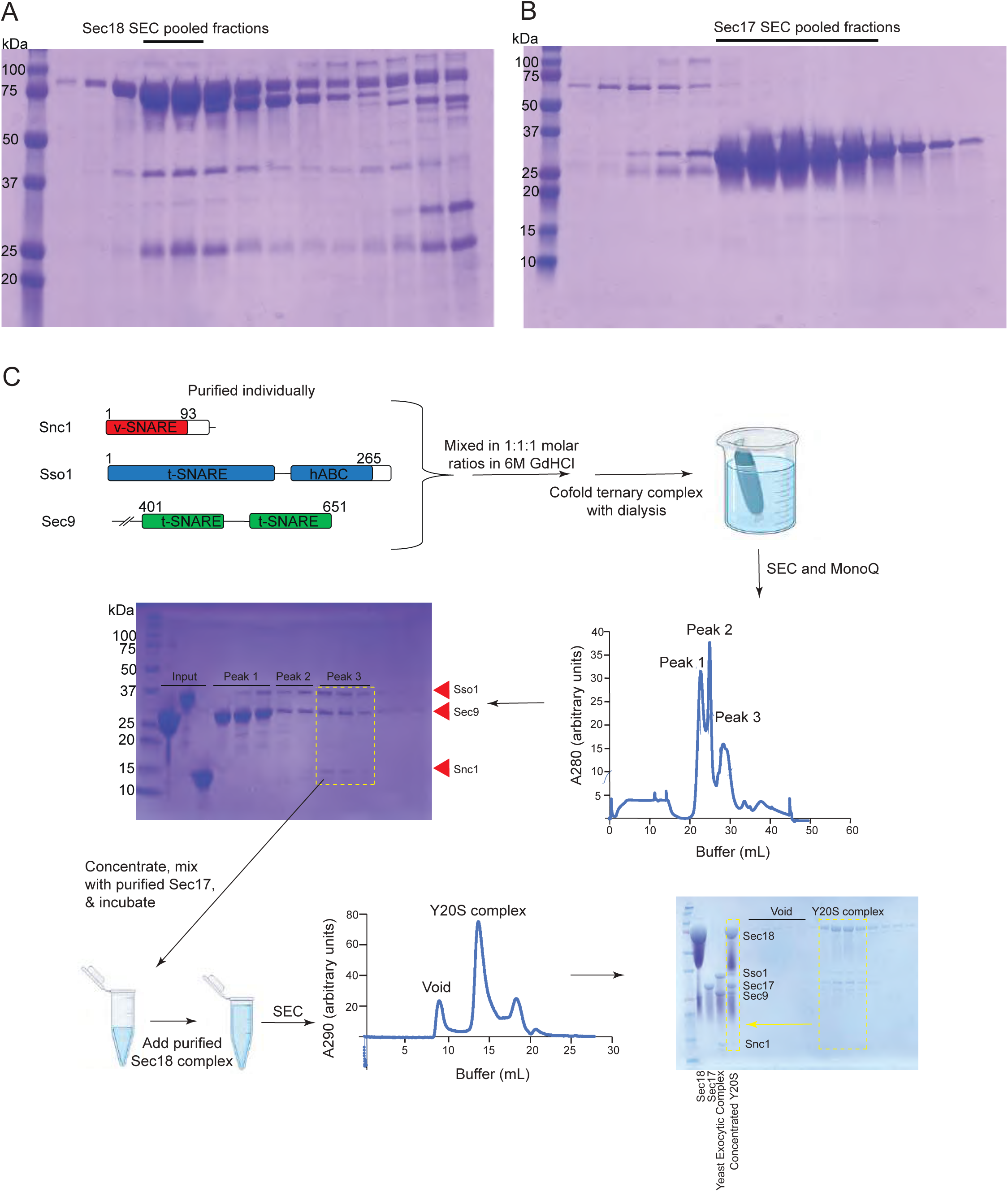

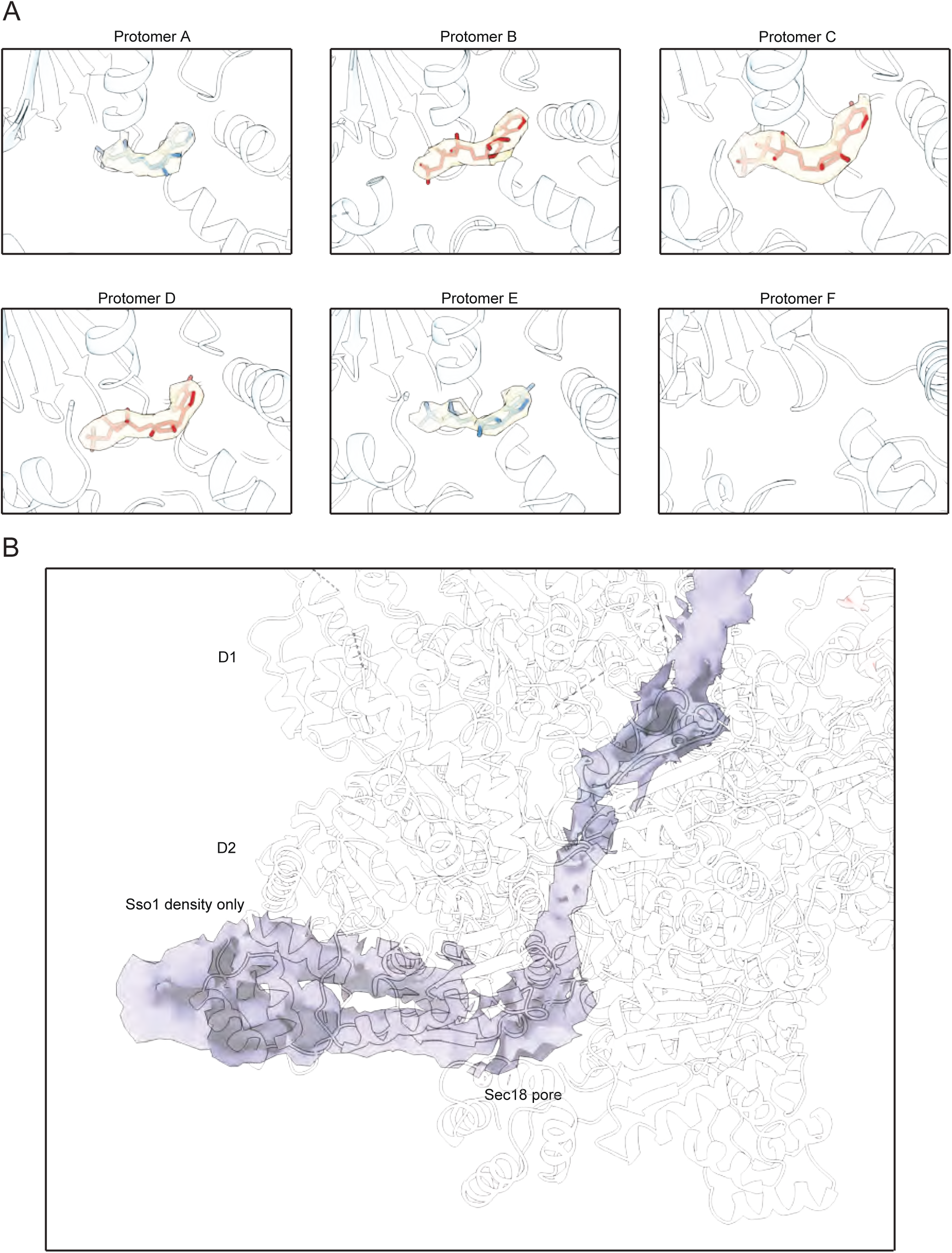

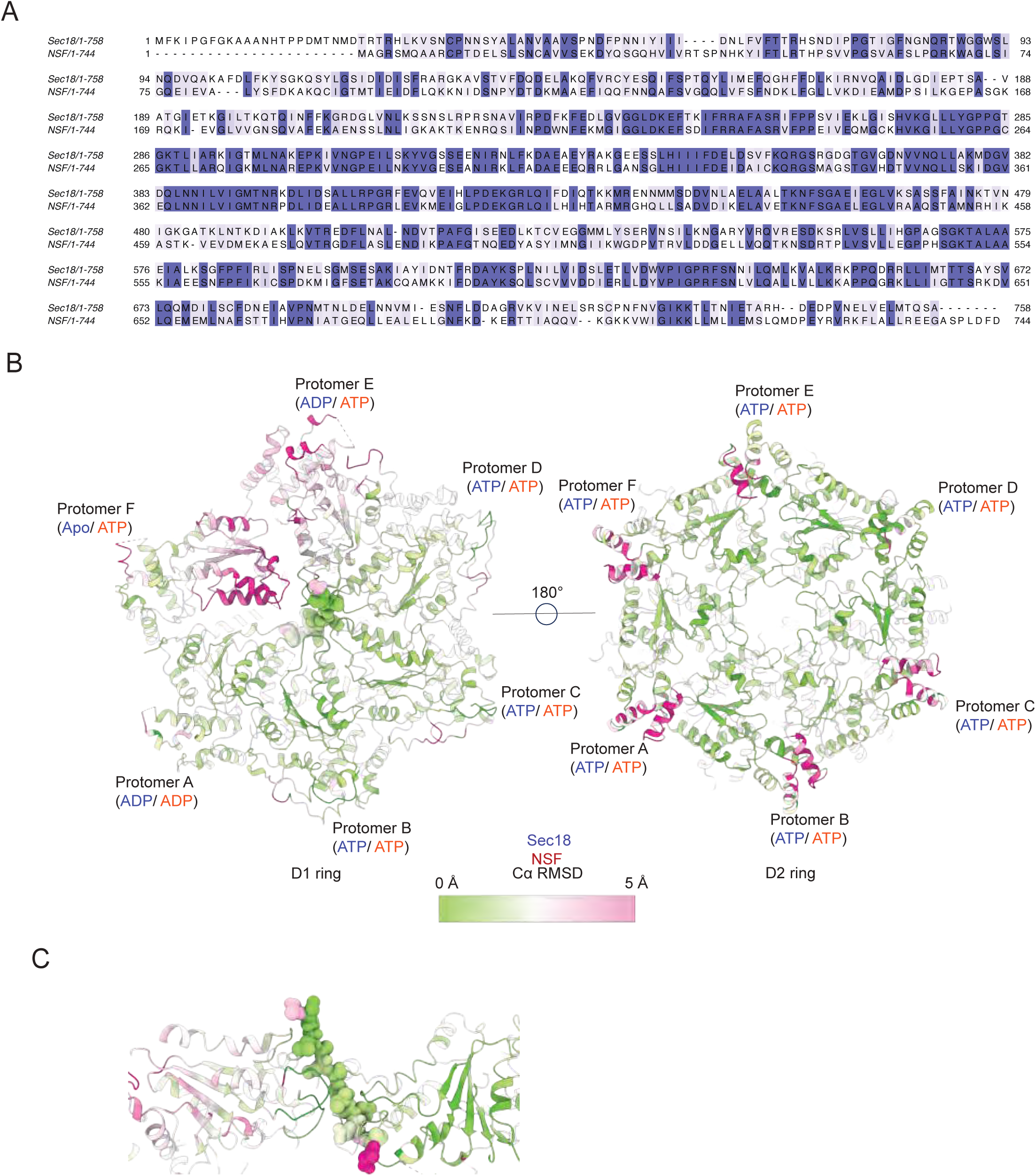

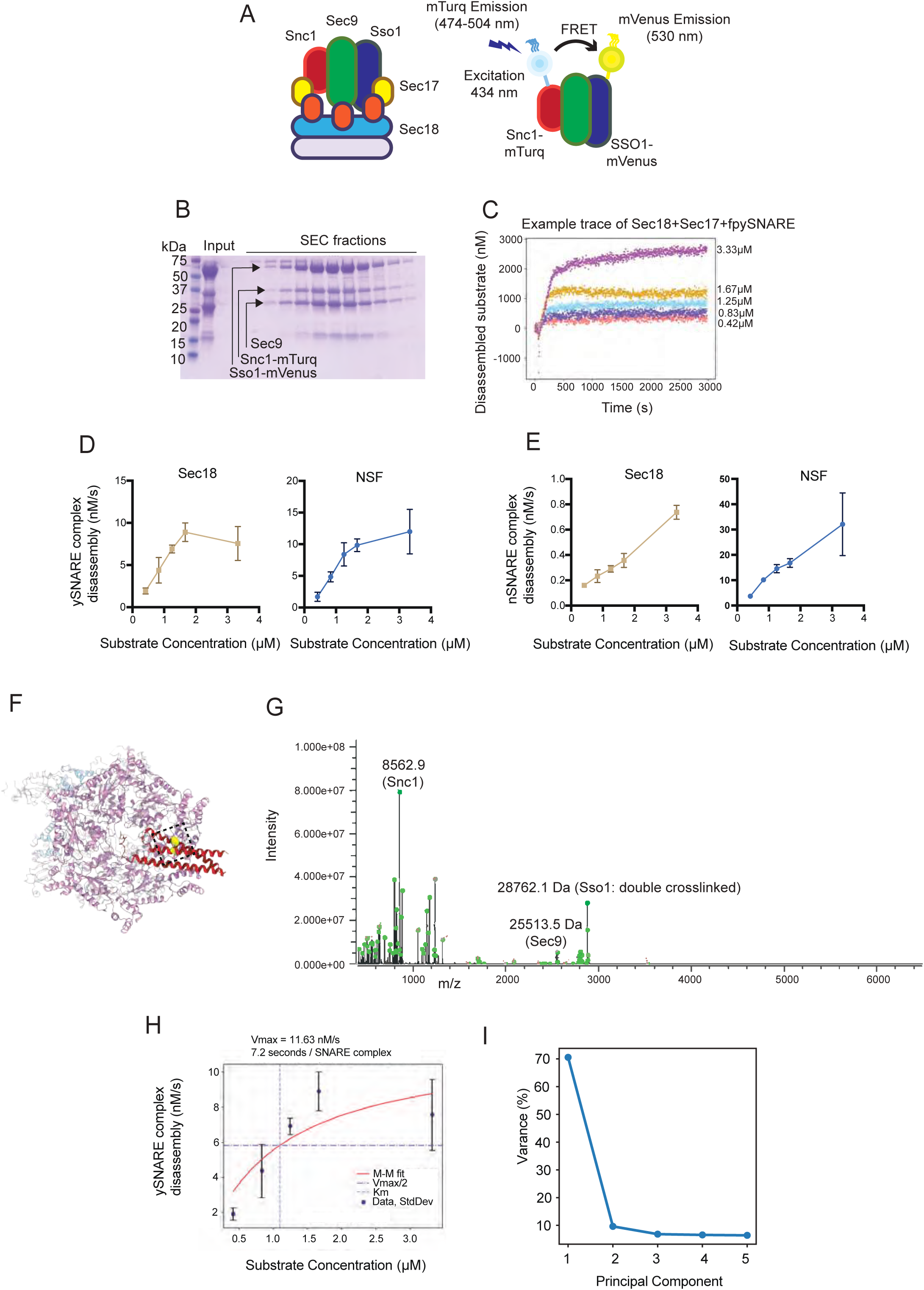

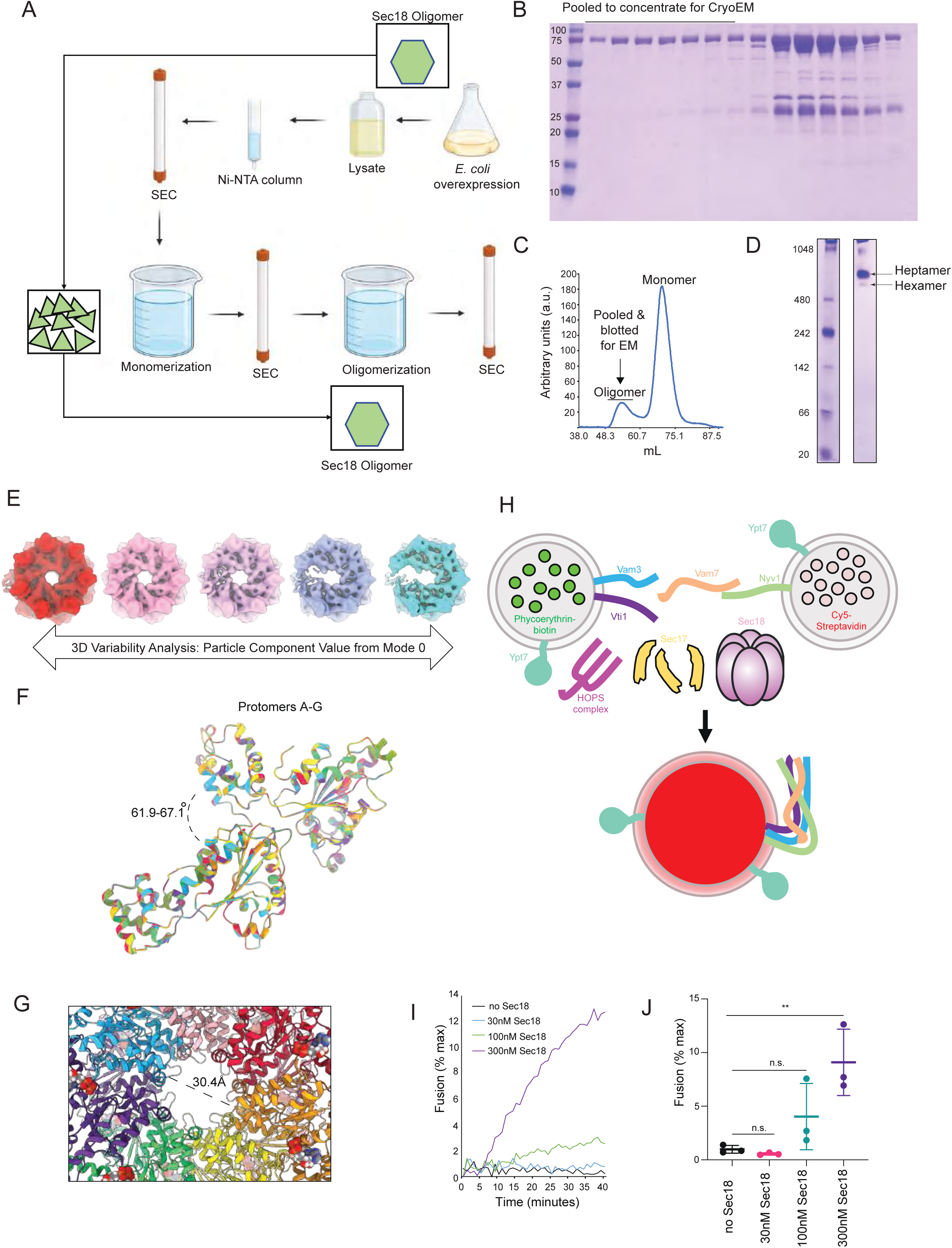

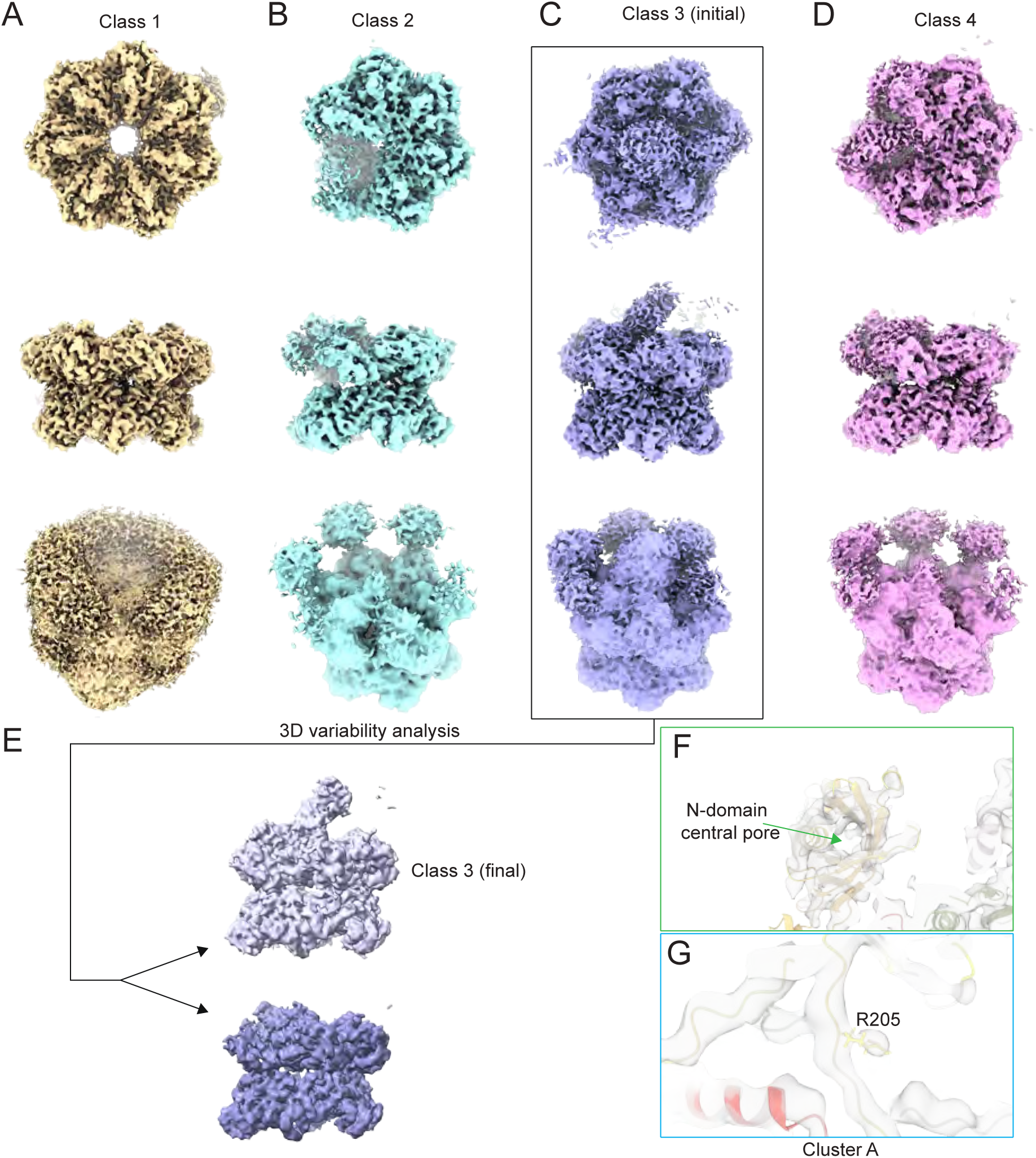

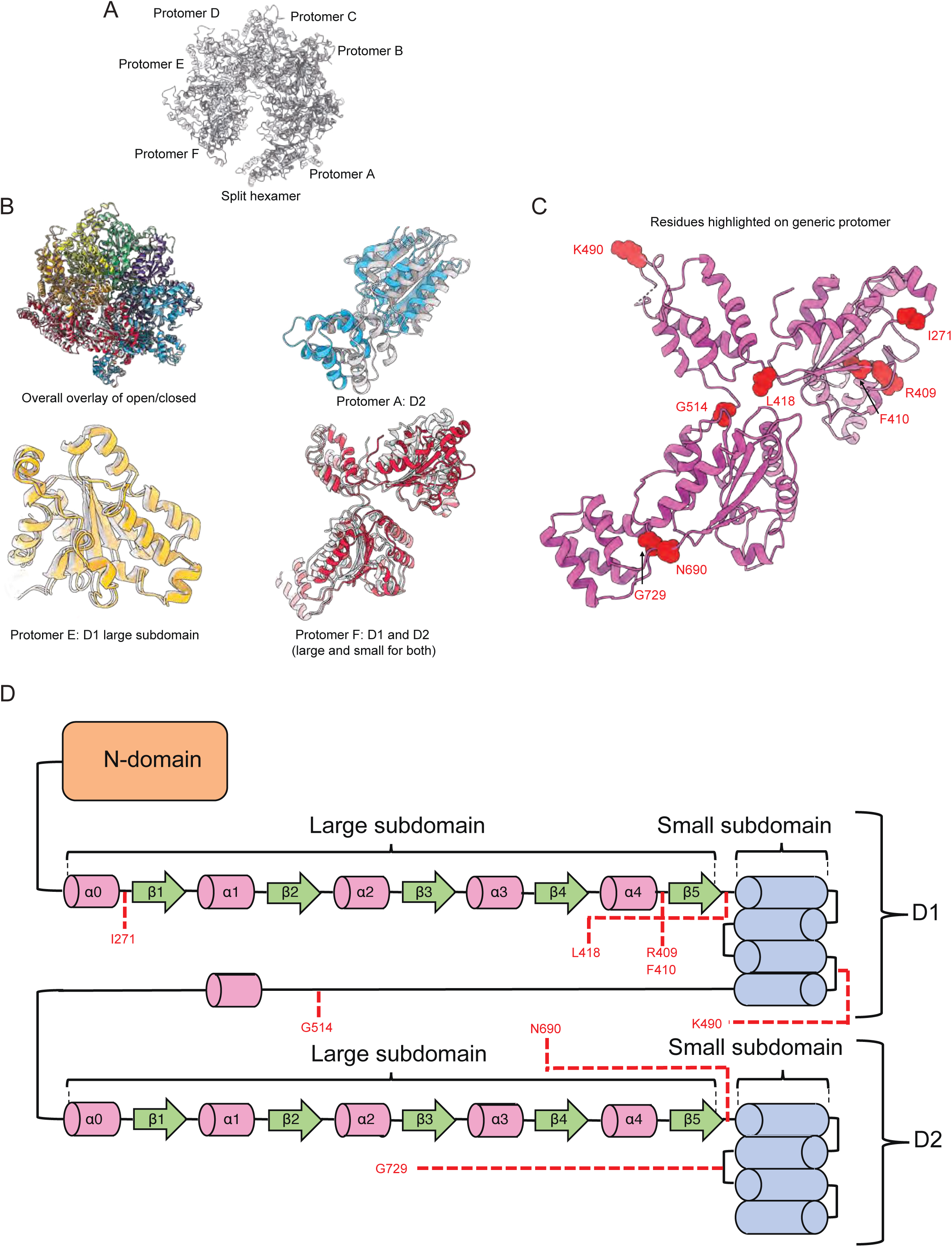

## Supplementary Information

### Supplementary discussion

#### Cross complementation experiments

In our cross-complementation experiments, Sec18 and Sec17 disassembled neuronal and yeast SNARE complexes (**Extended Data Figure 5D-E**), like NSF and α-SNAP disassembling ySNARE. NSF disassembled the ySNARE complex at a rate comparable to Sec18 (**Extended Data Figure 5D**), but, interestingly, Sec18 disassembled neuronal SNARE slower than NSF, ∼50-fold slower (**Extended Data Figure 5E**). This cross-complementation activity is corroborated by the high sequence and structural similarities between the yeast and neuronal systems. Despite differences in the SNARE substrate primary sequences, the substrate adopts a similar backbone conformation inside the pore of the D1 ATPase ring in both the yeast and neuronal systems (**Extended Data Figure 4C**).

#### Variability in N-domain engagement and Sec17 stoichiometry underlies substrate engagement

The eight classes from the non-hydrolyzing y20S dataset also show differences in the ySNARE-Sec17-Sec18-N-domain subcomplex arrangement (“spire”) and stoichiometry above the D1 ring of the y20S complex before disassembly (**Figure 2D-E**). Based on an unbiased Cα clustering algorithm (**Methods**), we found that the eight classes fall into four clusters. In cluster 1, classes 1 and 2 with two Sec17 adaptor molecules (the density for the third adaptor is weak) are bound by only three N-domains. In cluster 2, classes 3-5 with a stable, U-shaped formation of three Sec17 molecules are bound by 3 or 4 N-domains. Cluster 3 is defined by a more stable arrangement of the U-shaped Sec17 arrangement again but with an N-domain contacting at least one Sec17 adaptor protein. Lastly, in cluster 4, Y20S class 8 has all five proximal protomers’ N-domains engaging the Sec17 adaptor proteins.

Comparing the most extreme class averages in this dataset (class 1 and class 8, **Figure 2D**), the two Sec17 molecules lack the surface area to fully cover the exposed SNARE surface, which is covered upon binding a third Sec17 molecule (note that the linker between the two Sec9 SNAREs prevents a fourth Sec17 molecule from binding). This configuration also leads to a difference in the angle of the spire relative to the ring, where class 1’s spire has an acute angle relative to the D1 ring. In contrast, class 8’s spire is almost perpendicular. α-SNAP initially binds the SNARE complex with a 1:1 stoichiometry^7^, yet structural and single-molecule FRET data shows that the 20S can form with up to 4 α-SNAP molecules^8–11^. Forming a 1:1 complex of SNARE and adaptor allows for Sec18/NSF recognition via N-domain engagement. Then, additional Sec17 molecules bind to the spire before disassembly, allowing for additional connection points between Sec18 and the spire. Regardless of class, the arrangement of the protomers around the SNARE substrate in the D1 ring and the D2 ring arrangement is nearly identical, as the longer N-D1 linker is flexible enough to accommodate a variety of N-domain configurations.

As in the case of ɑ-SNAP, Sec17 interacts with the SNARE domain primarily through electrostatic interactions, consistent with structures of the neuronal 20S complex^8^. Alternating belts of electrostatic surface potentials complement each other in binding Sec17 adaptor proteins to the SNARE ternary complex (**Extended Data Figure 3B**). This observation may explain how Sec18 and Sec17 can process many SNARE complexes composed of different primary sequences. The N-domains and Sec17p domains have complementary electrostatic surface potentials, suggesting that charge-charge interactions play a role in this interaction. The N-domains form cavities of opposite charge that complement the C-terminal portion of Sec17 upon binding (**Extended Data Figure 3C**).

#### Vacuolar SNARE fusion assay

Vacuolar SNARE components were also found in our *in vivo* crosslinking experiment (**Figure 1B**), making the vacuolar an attractive candidate for a functional test. The vacuolar proteoliposome fusion assay mimics the homotypic fusion of lysosomes/vacuoles^12^ in which membranes expressing sets of SNAREs on each membrane must be preprocessed by Sec18, Sec17 and then passed to the HOPS complex to allow for homotypic fusion to occur. Using our preparation of Sec18, we observed fusion activity, which increased commensurately with the increase in Sec18 concentration. This observation confirmed that vacuolar SNARE processing was occurring, allowing homotypic fusion.

#### Substrate-free NSF under hydrolyzing conditions reveals additional conformations

We investigated whether NSF may also form these ring states under hydrolyzing conditions without an SNARE substrate. We purified NSF, exchanged it into a hydrolyzing buffer (*i.e.*, in the presence of Mg^2+)^, and determined cryo-EM structures. We found two primary oligomeric states, consistent with our Sec18 data, a flat heptameric (**Extended Data Figure 7A**) and a hexameric (**Extended Data Figure 7B-D**) state. These states of NSF are like the heptameric and split hexameric states of Sec18, with a flat D1 ring and ADP in all D1 protomers and ATP in all D2 protomers.

The D1 ring of hexameric NSF contains density in the pore (**Extended Data Figure 7B– E**); the most likely interpretation of this density is that NSF is engaged with part of the N-domain from one of the protomers (**Extended Data Figure 7F–G**). This structure suggests that Sec18/NSF tends to fill the side-loading gap in the absence of a SNARE substrate. Taken together, either Sec18/NSF accepts an extra protomer that can easily slide in or out (or binds to its N-domain), or it accepts a SNARE substrate, leading to hexameric 20S or Y20S complex formation. We note that we did not observe a pronounced coordinated ring opening in this condition, although starting from the assembled 20S complex, we did observe a coordinated split-open state of both D1 and D2 rings^18^. Instead, the particular conditions of the substrate-free NSF sample shift it to a self-inhibited hexameric state where the N-terminal domain is engaged with the D1 pore.

Our observations of heptameric or auto-engaged hexameric complexes, also seen in other AAA+ proteins^1,2^, follow a broader trend of inactive AAA+ conformations^3–6^ that appear reversible in the presence of substrate. We also note that the interaction between protomers differs depending on their oligomeric state (**Supplementary Figure 7**).

Whether or not Sec18/NSF heptamers are present in the cell at physiologically relevant conditions is unclear and will be subject to future investigations.

## Supplementary Figure Legends

**Supplementary Figure 1.**
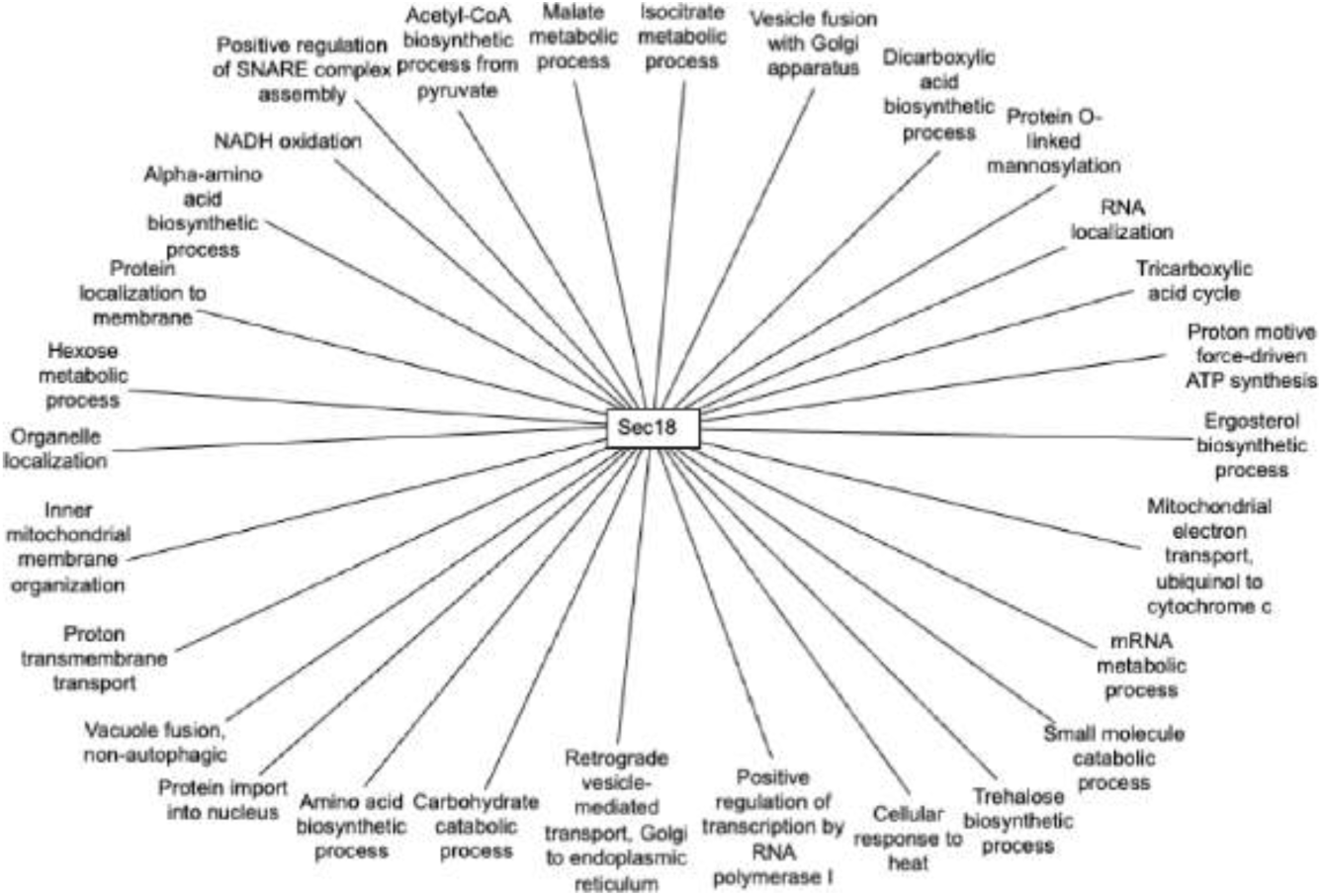
Gene ontology network of enriched processes based on enriched proteins in Sec18 coIP.

**Supplementary Figure 2.**
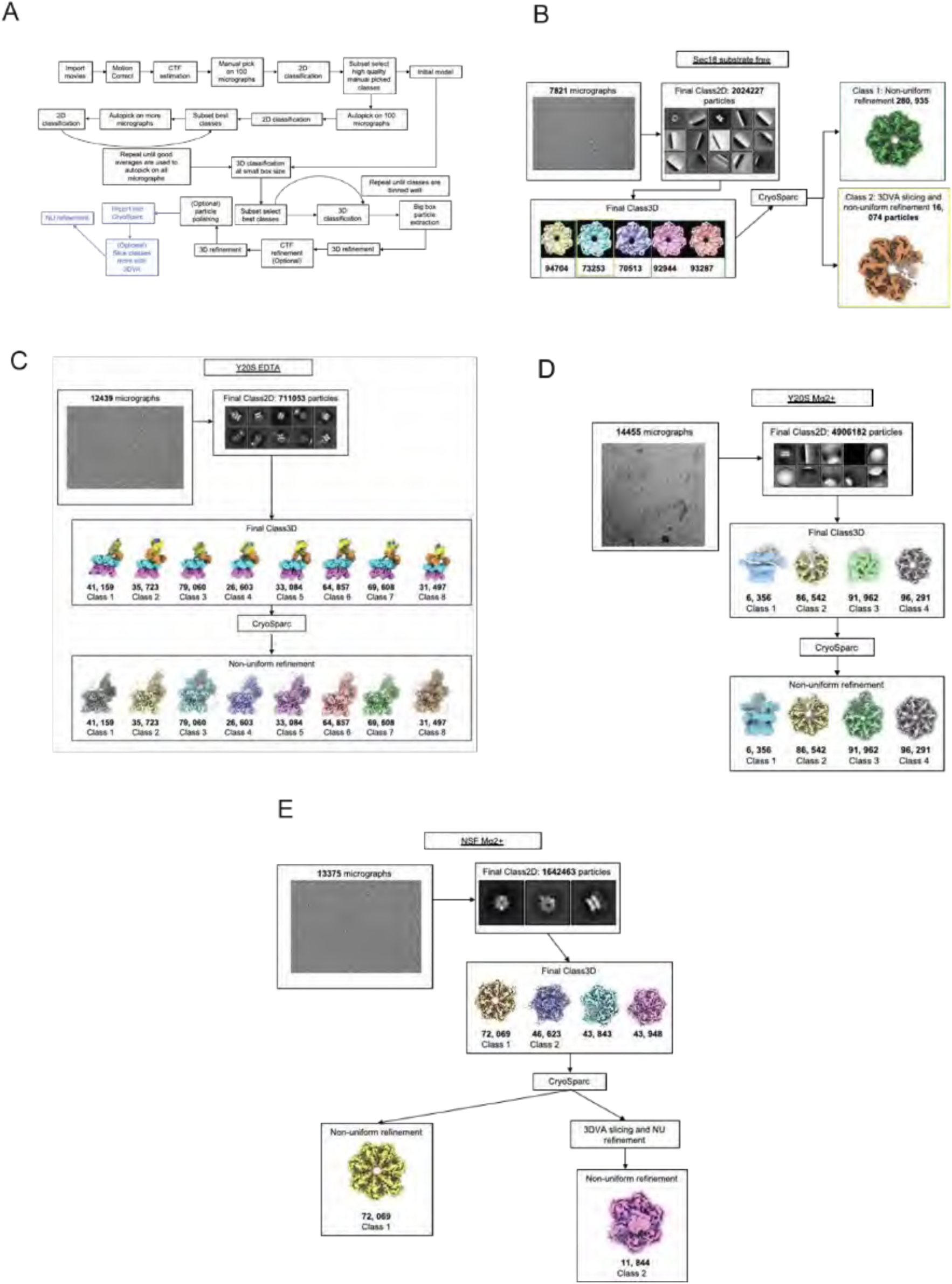
Processing information for the cryo-EM datasets. a) The general processing workflow that was employed for each dataset. b) Sec18 substratefree representative micrograph, Class2D, Class3D images, and final refinement depictions. c) Y20S EDTA representative micrograph, Class2D, Class3D images, and final refinement depictions. d) Y20S hydrolyzing condition representative micrograph, Class2D, Class3D images, and final refinement depictions. e) NSF hydrolyzing condition representative micrograph, Class2D, Class3D images, and final refinement depictions.

**Supplementary Figure 3.**
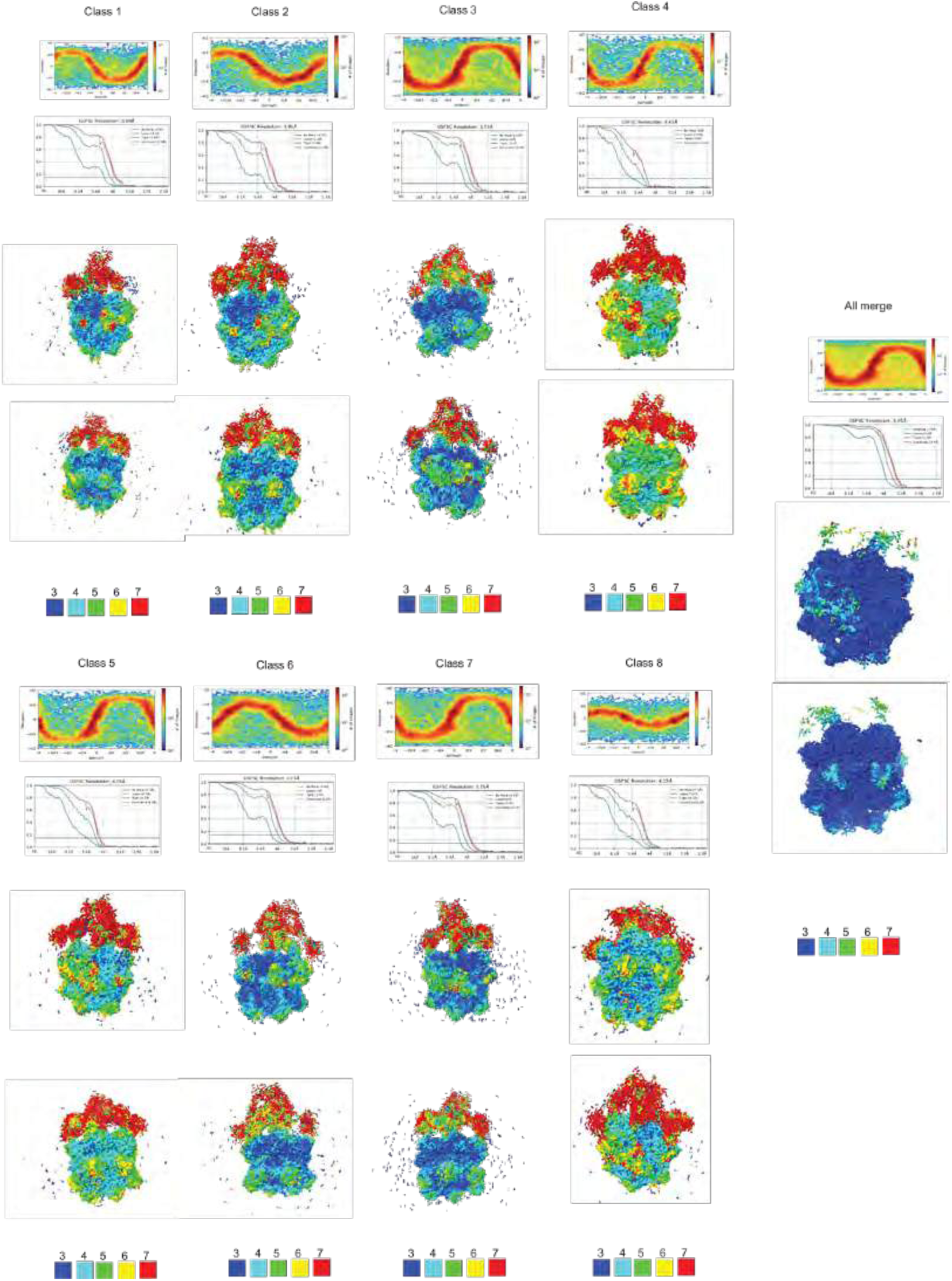
Distribution of particle orientation, FSC curves, and local resolution for all Y20S EDTA classes.

**Supplementary Figure 4.**
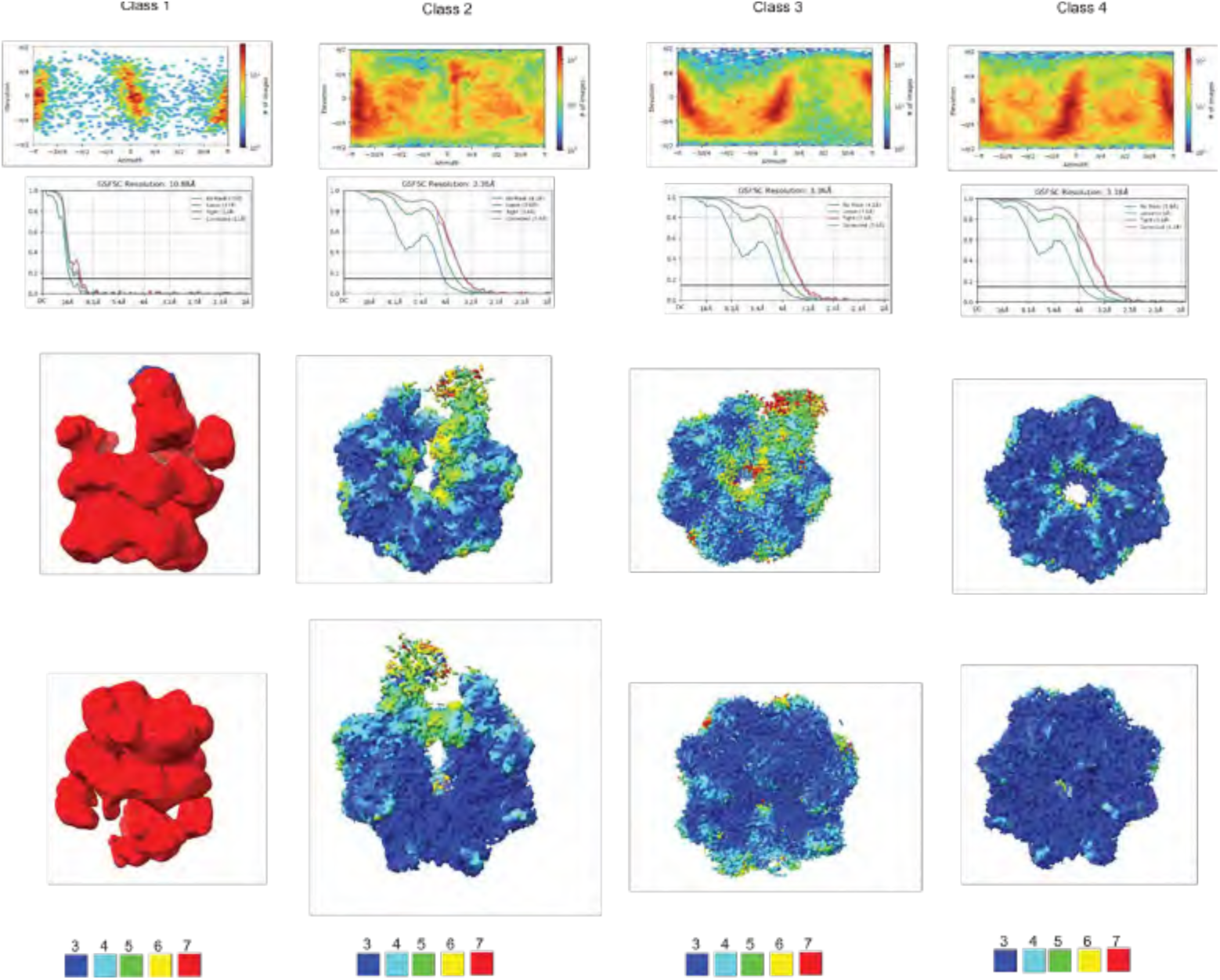
Distribution of particle orientation, FSC curves, and local resolution for all Y20S hydrolyzing classes.

**Supplementary Figure 5.**
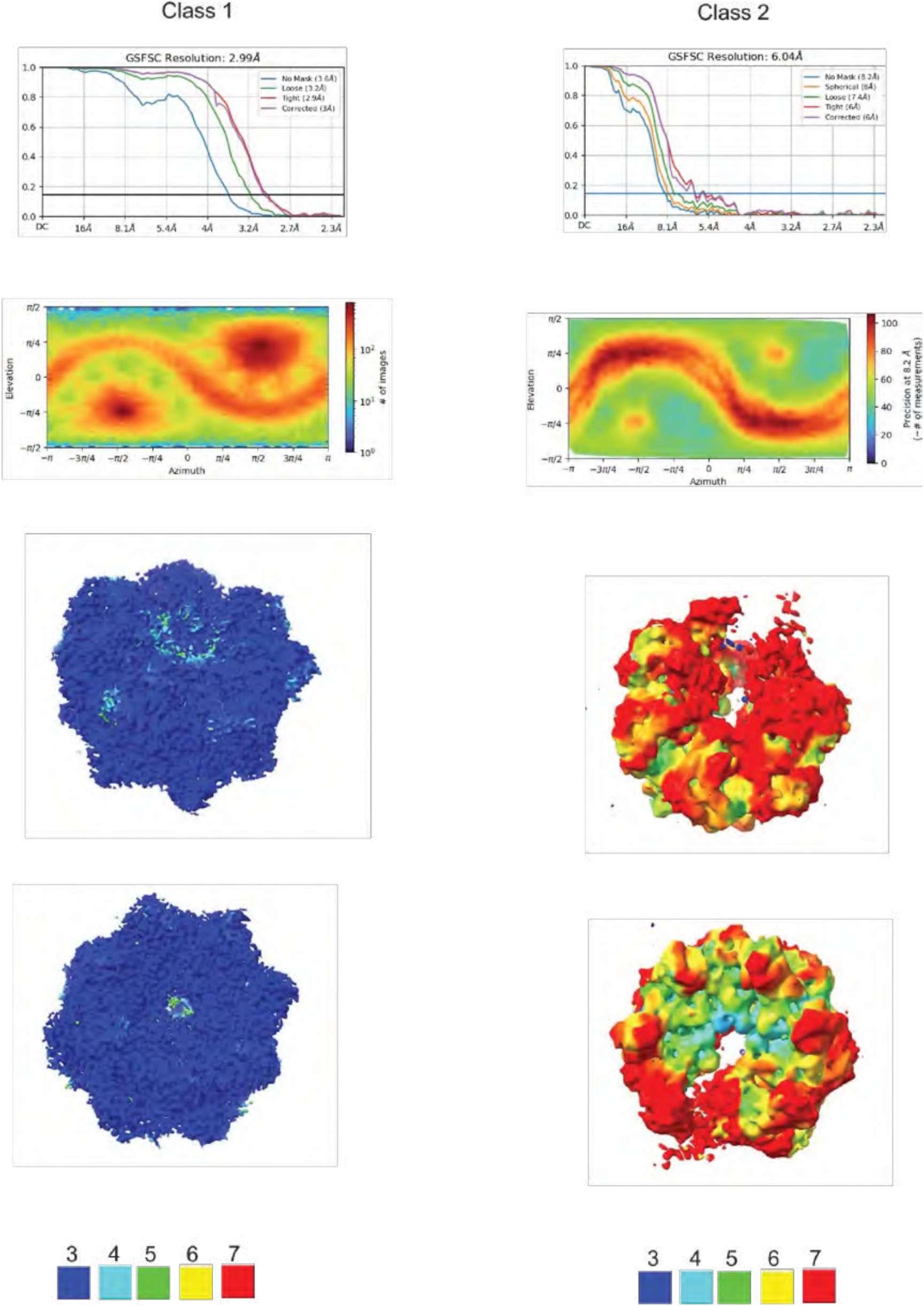
Distribution of particle orientation, FSC curves, and local resolution for all Sec18 hydrolyzing classes.

**Supplementary Figure 6.**
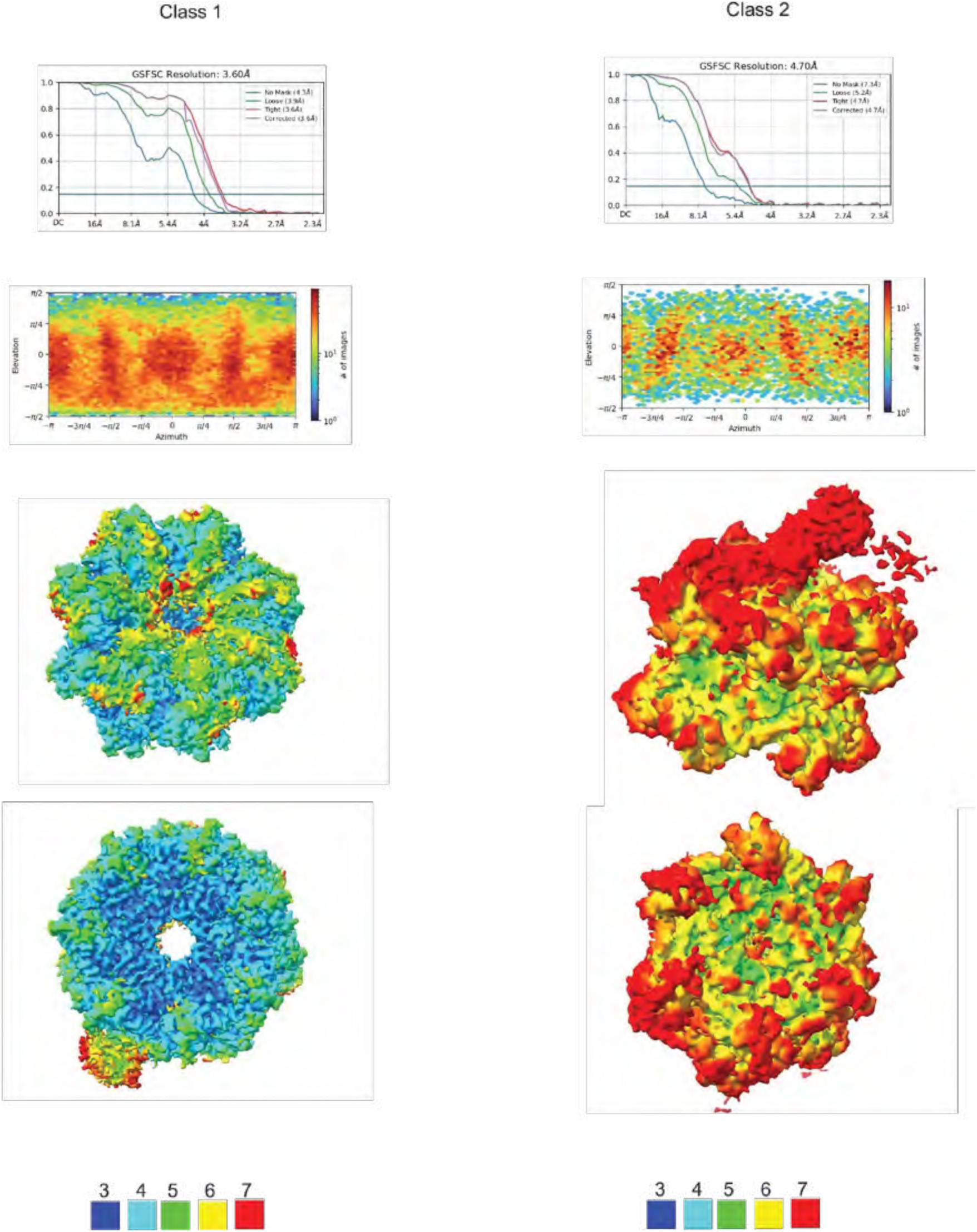
Distribution of particle orientation, FSC curves, and local resolution for all NSF hydrolyzing classes.

**Supplementary Figure 7.**
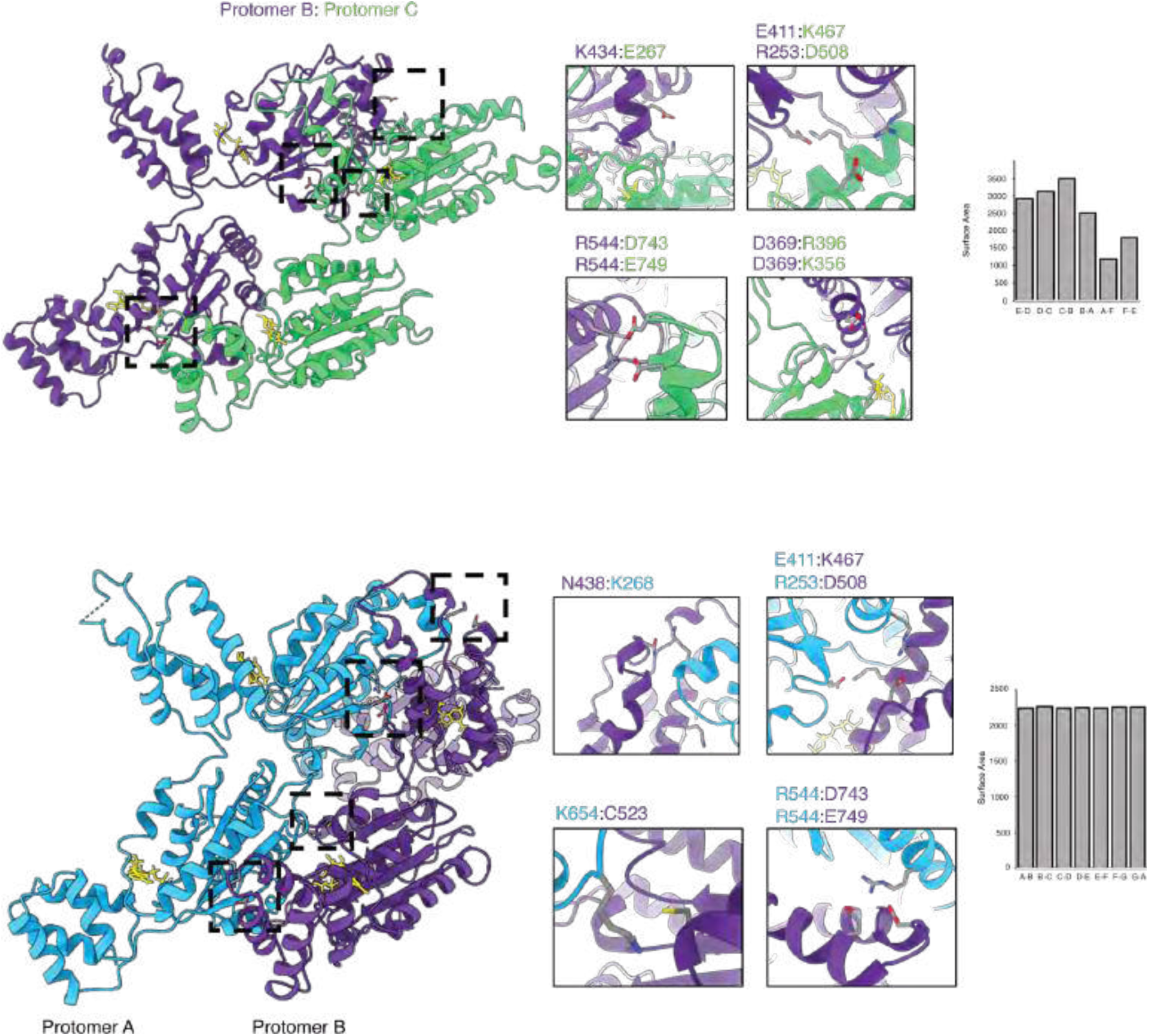
Interprotomer interface comparison of substrate-engaged (top) and substrate-free (bottom) protomers of Sec18. Interacting residues at the interface are shown.

**Supplementary Figure 8.**
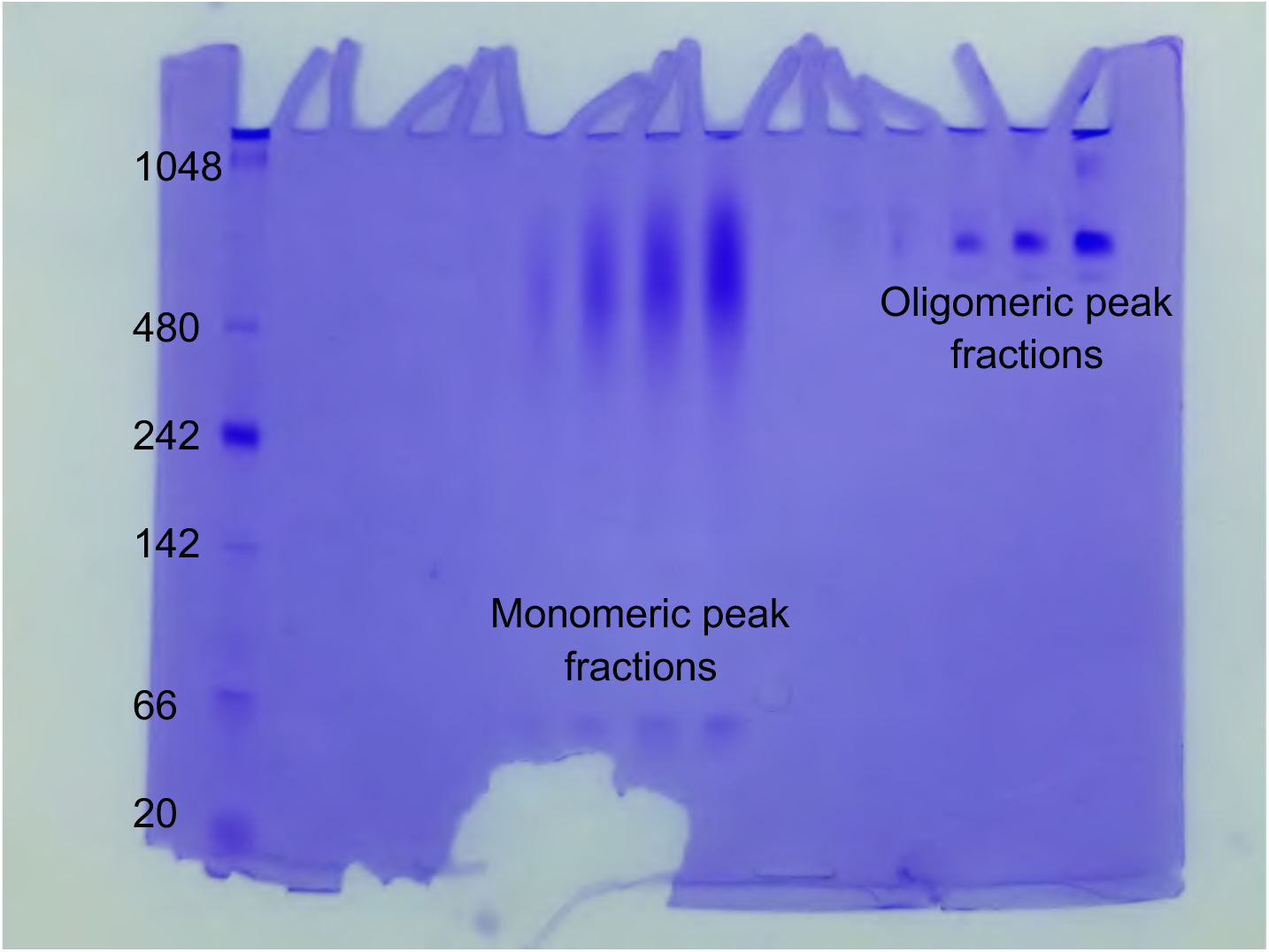
Full gel image of native gel of Sec18 hydrolyzing no substrate cryoEM sample, corresponding to cropped lanes shown in Extended Data Fig. 6d.

**Extended Data Table 1.**
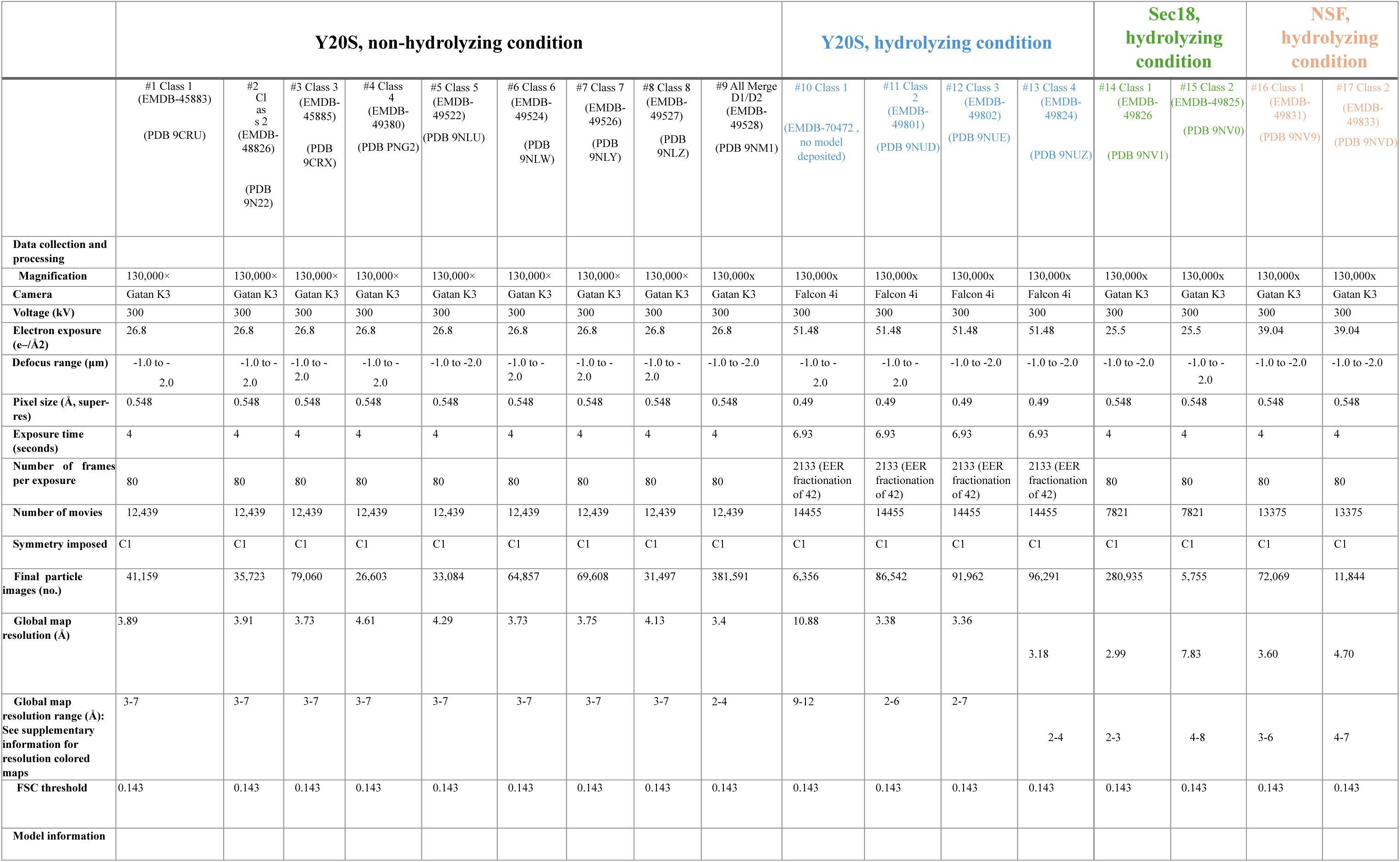
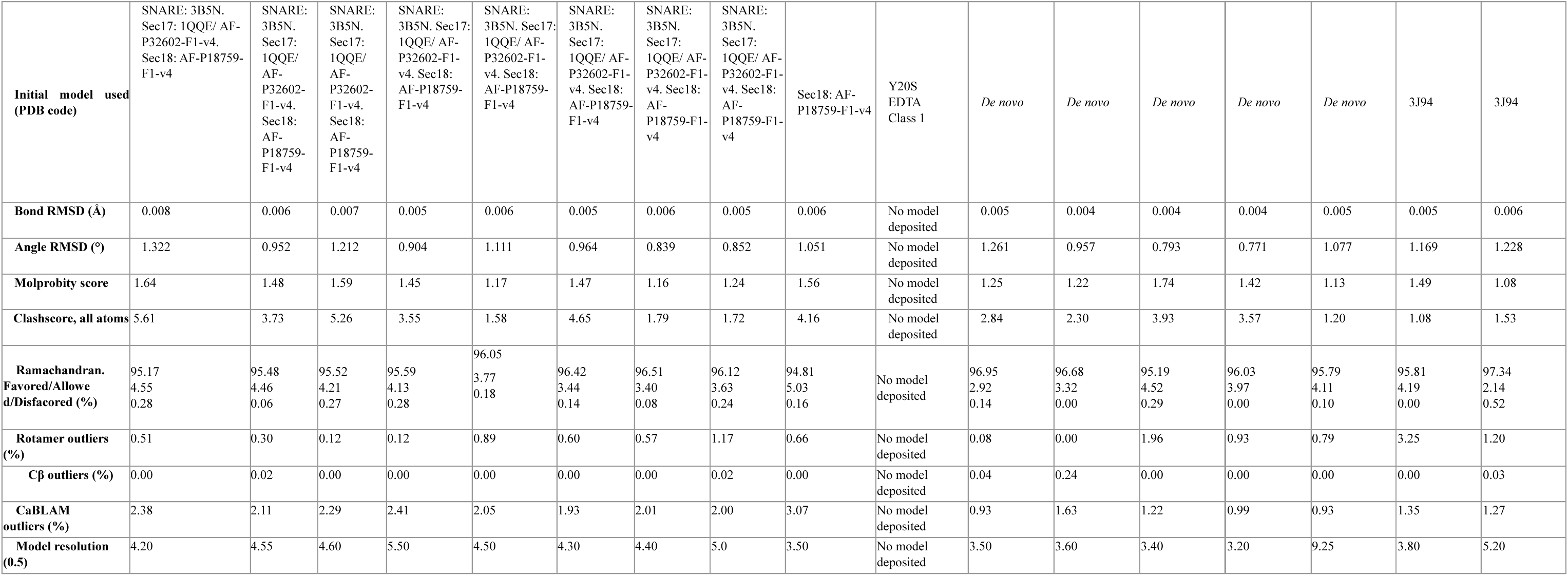
Cryo-EM data collection, refinement, and validation statistics

